# Interpretable EEG biomarkers for neurological disease models in mice using bag-of-waves classifiers

**DOI:** 10.1101/2025.08.14.670397

**Authors:** Maria Isabel Cano Achuri, Montana Kay Lara, Khalil Abed Rabbo, Benjamin T. Wilson, Austin Meek, J. Matthew Mahoney, Amanda E. Hernan, Austin J. Brockmeier

## Abstract

Electroencephalograms (EEGs) are time-series records of the electrical potential from collective neural activity in the brain. EEG waveform patterns—rhythmic and irregular oscillations and transient patterns of sharp waves or spikes—are potential phenotypical biomarkers, reflecting genotype-specific neural activity. This is especially relevant to diagnosing epilepsy without direct seizure observations, which is common in clinical settings, as well as in animal models, which often have subtle neurological phenotypes without overt epilepsy. Herein, we investigate genotypic prediction from long-term EEG signals of freely behaving mice belonging to six groups defined by the presence or absence of a neurological disease-genotype (*TSC1* gene knockout) in three different inbred strains with distinct genetic backgrounds. The potential complexity of genotype-related EEG patterns motivates a machine learning approach to automatically extract time-series descriptors, such as waveforms or spectral content, as biomarkers. We propose a machine learning approach to predict the genotypes of individual mice from the occurrence counts of waveforms that approximate short windows of the EEG. That is, a dictionary of waveforms is optimized to approximate windows from each genotype, and the vectors of waveform occurrence counts are the features for predicting genotypes via logistic regression models. Across two-fold cross-validation of the waveform dictionary learning, and leave-one-individual-out genotype prediction, we find that waveform counts pooled over multiple hour segments enable reliable prediction of mouse strain with an accuracy of 70% (chance rate of 38%), and for two of the three strains, DBA2 and C57B6, strain-specific classifiers reliably determined the epilepsy-genotype (*TSC1* gene knockout) at a 67% sensitivity with a 100% specificity for DBA2 and 67% specificity for C57B6. None of the mice of these strains had evidence of overt seizures or EEG-based seizure detection. The methodologies and results show the potential of EEG waveforms as phenotypes and bag-of-waves as a feature representation for identifying epilepsy genotypes.

## 1 Introduction

Electroencephalography (EEG) measures the superposition of electrical fields created by neural activity across the brain via electrodes placed at the scalp, cranium, or intracranially (iEEG) on the dura or on the cortical surface [1]. The dynamics in an EEG signal stem from the coordinated activity of populations of neurons, structurally and functionally networked, at time resolutions on the order of milliseconds [2]. The signal contains rhythmic oscillations, with power concentrated in well-known frequency bands, and transient waveforms [3, 4, 5]. These dynamics can be markedly disrupted in neurological disorders, such as epilepsy [6], which impacts 50–60 million people worldwide [7], due to underlying abnormalities at various scales from neural network structure to neuronal function. In cases of inherited diseases, we hypothesize that EEG biomarkers (descriptors of waveform patterns or frequency content) associated with genotypes can be useful to understand the link between genetics and neural network dysfunction.

We test this hypothesis in the case of a genetic panel of mouse models with a tuberous sclerosis complex (TSC)- associated mutation. TSC is defined by hallmark brain lesions that are often co-located with epigenetic foci [8]. TSC is caused by loss of function mutations in either the *TSC1* or *TSC2* genes, the products of which form a heterodimer that suppresses mammalian target of rapamycin (mTOR) signaling, which disrupts neural development [9, 10]. Even in carriers of TSC-causing variants without lesions or overt seizures, the primary effect of *TSC1/2* may be on neuron function that has wide spread effects on neural activity and behavior [10, 11]. Thus, we expect variation in brain function between knockout and wild-type across different genetic backgrounds. In the case of mouse models, the genetic background corresponds to inbred strains, which may have genetic modifiers that cause varying susceptibility to seizures in presence of *TSC1* knockout.

Here, we consider the problem of predicting the *TSC1*-genotype (haploinsufficiency caused by presence of *TSC1* knockout) from EEG biomarkers in conjunction with the background strain from time-series descriptors extracted from long-term, single-channel EEG recordings. We cast this biomarker discovery for epilepsy genotypes from long-term EEG recordings as a time-series classification problem. In this case, the true class of each EEG time-series is based on the genotype of the individual, and the goal is to model the relationship between EEG patterns and the genotype. We assume no knowledge of stimuli or behavioral activity.

The complexity of the patterns in EEG motivates using machine learning to automatically extract features and derive predictive biomarkers, as opposed to using predefined time-series descriptors or features. Crucially, these descriptors should be neuro-scientifically interpretable, which is often not the case for machine learning approaches. To extract interpretable features and biomarkers, we propose a machine learning approach that extends previous work on waveform learning for neural signals. This field was pioneered by foundational work using shift-invariant (convolutional) sparse coding to learn a dictionary of waveforms from single-channel time-series [12, 13, 14, 15, 16, 17], which can enhance later classification of the signals [18] and learn the morphology of non-sinusoidal brain oscillations [19, 20, 21]. To capture dependencies across EEG sensors, multivariate convolutional dictionary learning approaches have been developed [22, 23]. Recent work [24] has demonstrated the utility of these methods in identifying interpretable, transient neural events whose occurrence rates are biomarkers of aging. This highlights the potential of using learned waveform patterns as the basis for biomarker discovery, a principle we adapt in our present work.

Following recent work [25], we simplify the convolutional dictionary learning problem by working with short-time windows, assuming only one waveform is present in each window, and aggregate the number of times each waveform occurs across a longer segment. That is we do not consider the temporal dependence among windows, only within windows, which we term a ‘bag of waves’. To learn the waveform dictionary, we first cluster short windows from a single EEG channel from each genotype using a shift-invariant clustering algorithm. Each window in the training set is assigned to the nearest ‘centroid’ waveform, allowing for amplitude scaling and temporal shifts to best align. After all windows are assigned, the centroids are updated as the average of the windows after shifting to a common alignment. The set of centroid waveforms for a genotype acts as a dictionary. The vector of occurrence rates of each centroid waveforms in each genotype dictionary are concatenated to form the feature vector for predicting the genotype. This is analogous to the bag-of-words approach [26] used in information retrieval and document classification, and adapted to image/video retrieval [27], where instead of the count of words in a document, the waveforms are counted across a segment. As noted, the bag-of-waves feature representation does not carry information about the order of occurrence. After standard inverse frequency weighting that up-weights rare waveforms [28] and *ℓ*_2_-normalization, multinomial logistic regression (a linear model) is used to provide a calibrated probability for each genotype.

While the bag-of-waves features are based on counts of waveforms that are learned for each fold and genotype, many of the waveforms may not be useful for distinguishing the genotypes. For instance, waveforms learned for different genotypes may actually be similar in morphology and have occurrence rates across the genotypes. To identify the waveforms most critical to recognizing a particular genotype, we use Shapley values [29], which are simple to compute for linear models with an independence assumption: the Shapley value is simply the product of the coefficient and the difference of the feature value from its mean [30, 31]. We propose to find the waveforms’ Shapley values that correlate with a class of genotypes. Importantly, these could be waveforms with a negative coefficient for a particular class such that lower occurrence rate indicates a class. Thus, we examine waveforms with positive coefficients and Shapley values that correlate with a genotype. Together, the waveform shape, spectral content, and occurrence rate enable direct interpretation, leading to an understanding of each phenotype.

## 2 Data Collection

All animal procedures were approved by the University of Vermont IACUC and conducted in accordance with the Guide for the Humane Use and Care of Laboratory Animals. Mice were housed on a 12-hour light/dark cycle with ad libitum access to food and water, and behavioral testing was conducted during the light phase.

To leverage the advantages of a pure B6 background, mice used in this study were generated by first performing a backcross by breeding the commercially available conditional floxed *TSC1* mouse (*TSC1*^tm1Djk^/J, [32]; The Jackson Laboratories Strain #:005680), originally developed on a mixed B6, BALB/cJ, and 129/SvJae background, with pure B6 mice. In the first generation, female floxed carriers were crossed with male B6 mice to fix the Y chromosome; subsequent generations used male carriers and female B6 mice to fix the X chromosome. At each generation, we used The Jackson Laboratory’s Genome Scanning Service to assess background composition and selected progeny with the highest B6 content, accelerating the backcrossing process. By generation N5, we achieved over 99% B6 purity. This mouse is now available as Strain #:038428 at The Jackson Labs.

To generate a germline *TSC1* haploinsufficient mouse, we bred our pure B6 conditional *TSC1* floxed line (Strain #:038428) with a B6 mouse expressing Cre recombinase under the human CMV promoter (Strain #:006054), which drives ubiquitous expression. The resulting offspring had one *TSC1* allele knocked out in all tissues and one intact. This model enables breeding with genetic reference populations to systematically investigate how genetic background influences *TSC1*-related phenotypes. As genetic background strains, we include BXD87/RwwJ (BXD87, Strain #:007130) and both BXD reference population parent strains (B6, Strain #:000664 and D2, Strain #:000671). The resulting experimental mice—BXD87B6F1-*TSC1*^+/-^ (BXD87-Het), B6B6F1-*TSC1*^+/-^ (C57B6-Het), D2B6F1-*TSC1*^+/-^ (DBA2-Het)—were heterozygous for the *TSC1* knockout and inherited half the genetic background of the reference strain parent. Littermate controls (C57B6-WT, DBA2-WT, BXD87-WT) shared the same genetic background but lacked the *TSC1* knockout. Genotyping was performed on tail and toe snips collected prior to weaning. Data were collected from 45 mice in the resultant six groups. Both sexes were used. For consistency, we refer to WT/Het as the TSC genotype, the background genotype as the strain, and the combination of the TSC genotype and strain as simply the genotype. Table 1 summarizes the sex and genotype of all mice.

**Table 1:** Number of mice across genotypes and sex (F/M).

| Strain | Het |  | WT |  | Het | WT | F | M | Total |
| --- | --- | --- | --- | --- | --- | --- | --- | --- | --- |
|  | F | M | F | M |  |  |  |  |  |
| BXD87 | 7 | 3 | 2 | 5 | 10 | 7 | 9 | 8 | 17 |
| DBA2 | 4 | 2 | 3 | 5 | 6 | 8 | 7 | 7 | 14 |
| C57B6 | 4 | 2 | 6 | 2 | 6 | 8 | 10 | 4 | 14 |
|  | 15 | 7 | 11 | 12 | 22 | 23 | 26 | 19 | 45 |

### 2.1 Electrophysiology Recording

Electrodes were constructed with 6-pin Mill-Max strips attached to copper-tin wire leads. At approximately postnatal day 65, adult mice underwent implantation surgery. Mice were anesthetized with 1–2% isoflurane in oxygen and continually monitored for breathing rate and pain response. An incision was made at the top of the head to reveal lambda and bregma sutures of the skull. A needle was used to make six small burr holes in the skull where we placed skull screws (stainless steel). Copper-tin wire leads from the electrode were soldered onto the skull screws. The implant was sealed in place and covered to prevent discomfort or accidental removal. The six pin holes were exposed at the top. After the electrode was securely attached to the head, the mouse recovered for five days. After recovery, we attached a Pinnacle headmount (Pinnacle Technologies, Lawrence, KS) to the electrode implant and recorded EEG using Pinnacle data acquisition software that extracts live data via Bluetooth and concurrent video monitoring.

Signals were high-pass filtered at 1 Hz and sampled at a rate at 256 Hz. Three channels were recorded: 1) a right frontal signal relative to the reference signal, 2) a left frontal signal with the same reference, and 3) a bimodal parietal signal. Recording lasted five to seven days and the mouse was allowed to freely move, eat, and sleep in an open cage. Table 2 details the statistics of the recording lengths by group.

**Table 2:** Statistics of the recording length (hours) across genotypes.

| Genotype | Count | Ave. | Min | Max | Total |
| --- | --- | --- | --- | --- | --- |
| BXD87 Het | 10 | 121.1 | 51 | 163 | 1211.4 |
| BXD87 WT | 7 | 106.9 | 52.4 | 140.4 | 748.6 |
| DBA2 Het | 6 | 137.4 | 70.6 | 228 | 824.6 |
| DBA2 WT | 8 | 119.2 | 87.6 | 134.2 | 953.6 |
| C57B6 Het | 6 | 123.2 | 100.6 | 143.2 | 739.2 |
| C57B6 WT | 8 | 109.7 | 94.2 | 131.8 | 877.6 |

Spontaneous seizures were identified electrographically by visual inspection and Sirrenia seizure detection software (Pinnacle Technologies, Lawrence, KS), and confirmed behaviorally with video monitoring offline (data not shown). Of the 45 mice, spontaneous seizures were identified in 3 mice of BXD87 Het genotype, with a total of 57 seizure events, averaging 5.12 minutes. Specifically, the first mouse had 10 events averaging 4 minutes, the second had 27 events averaging 6.46 minutes, and the third had 22 events averaging 5.15 minutes.

In subsequent analysis of the dataset, we only use the first channel (a right frontal referential signal) and in the three BXD87 mice with seizures we exclude portions of the recordings 5 minutes before the start and 60 minutes after the end of each event. Because of consecutive seizures the exclusion regions overlap, there are 10, 8, and 8 exclusion periods, averaging 2.15 hours, 4.15 hours, and 2.55 hours, and totaling 21.5, 33.3, and 20.4 hours for the three mice respectively.

### 2.2 Dataset Division

As shown in Table 3, the dataset is divided into two-folds by pseudo-randomly assigning each individual to the folds with equal odds. To check consistency of the entire genotype classification approach, five random two-fold splits are created, but the qualitative analysis will focus on the first split.

**Table 3:** Counts of individuals across genotypes, sex (F/M), and fold (0/1).

| Fold | Strain | Het |  | WT |  | Het | WT |
| --- | --- | --- | --- | --- | --- | --- | --- |
|  |  | F | M | F | M |  |  |
| 0 | BXD87 | 3 | 2 | 1 | 2 | 5 | 3 |
| 1 | BXD87 | 4 | 1 | 1 | 3 | 5 | 4 |
| 0 | DBA2 | 1 | 2 | 3 | 1 | 3 | 4 |
| 1 | DBA2 | 3 | 0 | 0 | 4 | 3 | 4 |
| 0 | C57B6 | 3 | 0 | 4 | 0 | 3 | 4 |
| 1 | C57B6 | 1 | 2 | 2 | 2 | 3 | 4 |

## 3 Machine Learning Methods

The proposed machine learning approach to solve the time-series classification task involves class-conditional feature learning followed by supervised classification. The features are derived from a bag-of-waves, which are counts of different waveforms organized into class-conditional dictionaries, occurring in segments of the EEG. A linear classifier in the form of logistic regression is then trained to predict the class label of each segment. To predict the class of an individual, a single bag-of-waves is calculated across segments sampled throughout the time series. Classifiers are trained to predict the joint genotype, components of the genotype (strain and TSC-genotype). Additionally, factorized classifiers for the joint genotype are constructed by first predicting the background strain, and then using a classifier for the TSC-genotype given the strain. Finally, feature analysis is conducted with Shapley values, using a simplified formula [30, 31], applicable to independent features for a linear model. The complete set of code is available at Anonymous GitHub.

### 3.1 Bag-of-Waves Representation

The bag-of-waves features are based on learning a dictionary of waveforms that can be used to approximate short windows of the EEG from each class. Each waveform is shorter than the window it seeks to explain and the goal is to find the waveform with the optimal shift and scaling that approximates the window. Figure 1 shows an example of a 1-second waveform best matched to a portion of a 2-second window.

**Figure 1:**
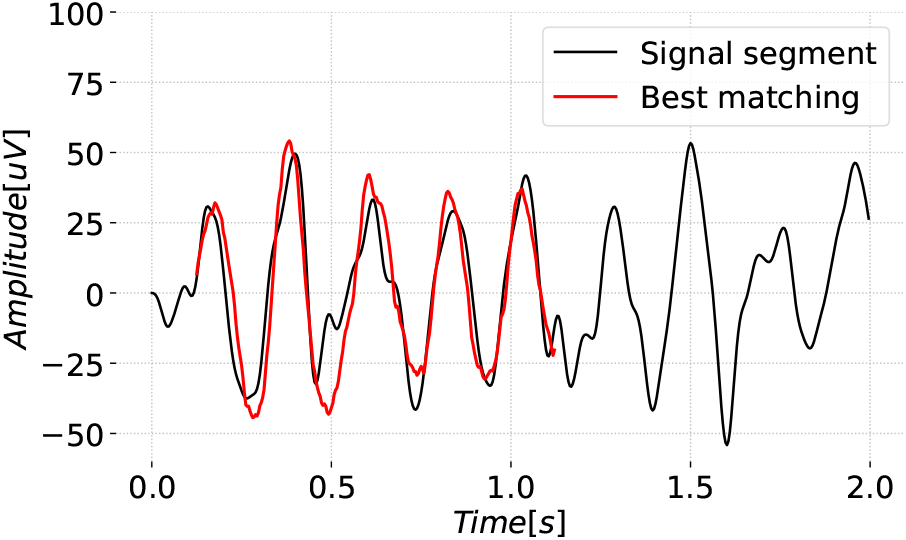
Best matching 1 second waveform for a 2 second segment is shown shifted and scaled.

The dictionary learning algorithm (shift-invariant k-means with cosine similarity) is applied to windows extracted from signals for all individuals in the training set. During learning, a single waveform from the current dictionary is matched in terms of cosine similarity to a portion of each window. Cosine similarity implicitly finds the optimal scale of the waveform and ignores the norm of the signal while matching. Then, each waveform in the dictionary is updated such that it is the average of the portions of windows for which it best aligns. Here, the goal is that each dictionary is tailored to capture the commonly occurring patterns within the EEG recordings for each genotype.

After dictionary learning, EEG signal segments are characterized by the bag-of-waves determined by the counts of waveforms best matched to windows. This is done for each of the class-specific dictionaries, and the counts of each waveform across all dictionaries are concatenated into a single feature vector. Waveforms of 1 s are matched to portions of non-overlapping 2 s windows taken from 1-hour EEG recording segments. While the bag-of-waves from 1 hour segments are used for training the classifier, to classify an individual, the counts of multiple segments can be pooled together, or shorter segments can be tested.

The individual dimensions of the vector can be normalized to down-weight overly frequent waveforms and up- weight rarer, possibly discriminative waveforms. For this purpose, we apply the inverse document-frequency (IDF) normalization [33, 28], where the document frequency refers to the number of segments that the waveform is nonzero, is estimated on training data. Finally, the weighted vector is normalized to unit-norm, and this is the feature representation for subsequent classification. The normalization ensures that the feature representation is the same whether counts or rate (counts per time unit) are used.

#### 3.1.1 Time-series Windowing and Shifts

Mathematically, we denote a discrete-time signal of length *L* as 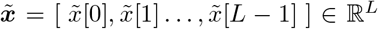, where 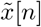 represents the value at discrete time index *n* ∈ {0, …, *L* − 1}. A *P* -length window of this signal starting at time point *τ* ∈ {0, …, *L* − *P*} is obtained by the window operator

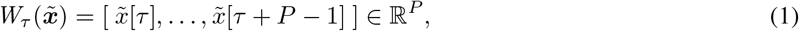

with *P < L*. Note that *W*_*τ*_ is a linear operator, 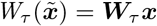, where ***W***_*τ*_ is binary matrix with *P* -nonzero entries, and its adjoint operator *S*_*τ*_ takes an *L*-length window 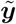 and pads it with zeros, shifting it to start at time point *τ* :

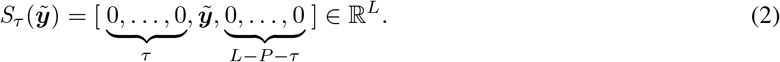

Note that 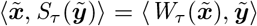, where ⟨***a, b***⟩ = ∑_*k*_ *a*_*k*_*b*_*k*_ denotes the dot/inner product.

#### 3.1.2 Shift-Invariant Dictionary Learning

Sparse coding is based on the assumption that signals (data vectors) from a specific random process (distribution) can be approximated as a combination of a relatively small number of waveforms (basis elements or atoms) chosen from a dictionary [34, 35]. The dictionary is specific to the process or distribution, and may be learned by trying to approximate signals with a sparsity constraint.

Here we consider dictionary learning and sparse coding in the convolutional case [12, 13, 14, 15, 18, 17, 16, 19, 20, 21], where waveforms can appear at any shift. Mathematically, a continuous-time signal 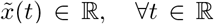 can be approximated as a sum over *A* atoms, where the *i*th atom consists of a waveform 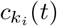 chosen from the dictionary of waveforms 𝒞 = {*c*_1_(*t*), …, *c*_*K*_(*t*)} scaled by *α*_*i*_ ∈ ℝ_≥0_ and shifted by 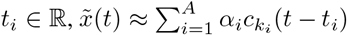. Practically, the time-series is sampled at a rate of *f*_*s*_ Hz (sampling interval of 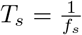), yielding a finite discrete time series 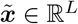 of length *L* (a duration of *LT*_*s*_). Then the shifts are rounded to whole numbers *τ*_*i*_ ≈ *t*_*i*_*f*_*s*_, and

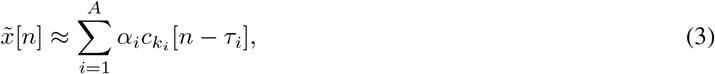

where ***C*** = [***c***_1_, …, ***c***_*k*_] ∈ ℝ^*P* ×*K*^ denotes the matrix corresponding to the discrete-time dictionary of waveforms of length *P, P < L*. Using the shift operator *S*, the vector form is

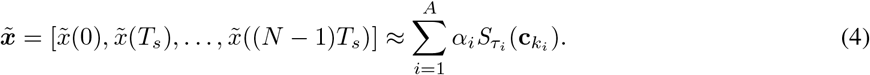

Given a dictionary and number of atoms *A*, the parameters of the atomic representation 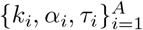 are typically optimized to minimize the sum of the squared reconstruction error. Approximate solutions can rely on convex relaxations such as basis pursuit [36] or greedy approaches such as matching pursuit [37] or orthogonal matching pursuit [38, 39]. As learning the dictionary requires repeatedly solving this problem while jointly optimizing the dictionary, it is even more computationally challenging [40, 41], especially in the shift-invariant case [19, 42, 17, 43]. To limit some of the computational challenges of dictionary learning in convolutional sparse coding, we consider an extreme case of sparsity where only a single waveform is dominant in each relatively short window. As each window will be assigned a waveform, this corresponds to a clustering problem.

#### 3.1.3 Shift-Invariant k-Means Clustering

Given *M* windows of length 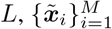, we use a shift-invariant k-means algorithm [25] to learn a dictionary of *K* waveforms ***C*** = [***c***_1_, …, ***c***_*K*_]^⊤^ ∈ ℝ^*K*×*P*^ of length *P*, ***c***_*k*_ = [*c*_*k*_(0), …, *c*_*k*_(*P* − 1)].

The algorithm is motivated by finding the dictionary of waveforms that best approximates the given windows, in terms of minimizing the mean squared error, by shifting and scaling one of the waveforms for each window, as in the following minimization problem:

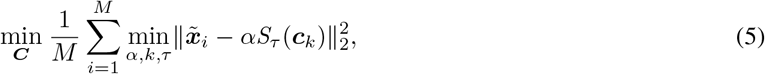

where 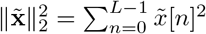 is the squared *ℓ*_2_-norm (Euclidean norm).

The shift-invariant k-means algorithm [25], is an alternating optimization, since given the dictionary each window can be separately matched to a waveform in terms of cosine similarity, and given the assignments and shifts the waveform can be updated as the average of the corresponding windows. To understand this approach, we expand the squared error for 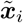

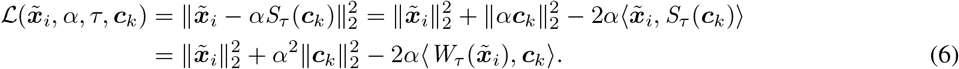

For given *τ* and *k*, the optimal *α* (ignoring the non-negativity) can be found by finding the root of the partial derivative, which yields 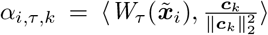 and 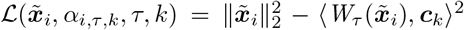. The inner optimization is then 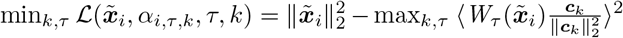. Because the sign of 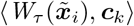 corresponds to the sign of *α*_*i,τ,k*_, the minimal loss with the non-negative coefficient is 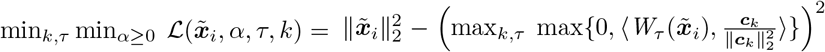, where *k, τ* can be chosen arbitrarily when all the coefficients are negative such that the second term is zero. Thus,

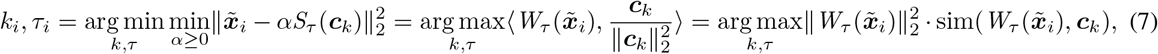

where 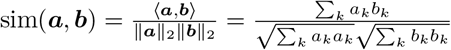 is the cosine similarity. Assuming window has constant magnitude in each *P* -length sub-window, 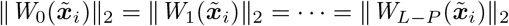, the inner optimization is

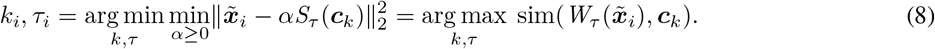

Generally, equation 8 does not hold, as the best assignment and shift for cosine similarity may not minimize the squared error since the cosine similarity ignores the norm of the sub-windows. Nonetheless, in synthetic simulations, where the true waveforms are known, we find that cosine similarity provides matching that yield waveforms that are more similar on average to the true waveforms, compared to equation 7.

Given the assignments, the goal is to update the waveforms to minimize 5. This can be broken into *k* subproblems by collecting the windows each waveform appears in ℐ_*k*_ = {∈ {*i* 1, …, *M*} : *k*_*i*_ = *k*}, *k*∈ {1, …, *K*}. Ignoring the sign constraint on the coefficients, the optimal waveform update is actually provided by the rank-1 truncated singular-value decomposition (SVD) of the matrix 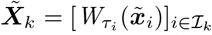 [40]. However, the SVD is not robust in the sense that the influence grows quadratically with the scale of one window; thus, a single outlier window can breakdown the estimate. This has motivated using *ℓ*_2_ normalization to each window before applying SVD [44, 45]. The results of our simulation also show that the averaging used by the shift-invariant k-means, works better than SVD or methods that incorporate the scaling [46]; however, averaging using *ℓ*_2_ normalized windows, which is optimal for cosine similarity 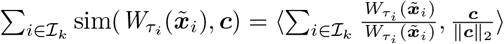 has a slight performance increase in simulation. Due to concerns that windows with low-norm would be noise in the real recordings, we did not adjust the algorithm.

The complete algorithm [25], starts from an initial dictionary ***C***^(0)^ and across iterations *m* = 1, 2, … performs three steps:

1. Assignment step:

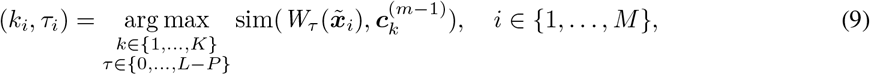
2. Window collection step:

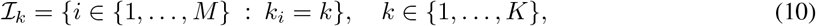
3. Waveform update step:

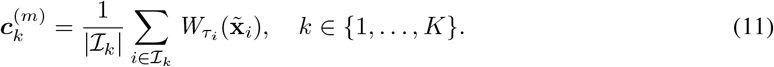

The optimization and this algorithm depart from the standard k-means algorithm by the search for the best shift and by using cosine similarity for the assignment step, which together allow the waveform to shift and scale to match the window. Like k-means, in the waveform update, a simple average is conducted. The complexity of each iteration of the algorithm is 𝒪 (*MKL*).

##### Specifics for our dataset

For the mouse-genotype data, we use *J* = 6 dictionaries ***C***_1_, …, ***C***_*J*_ for the six genotype groups, each with *K* = 200 waveforms of length *P* = 256 (1 second at the sampling frequency of 256 Hz) for windows of length *L* = 512 (2 seconds). Each dictionary is learned from *M* = 40, 000 windows drawn across all individuals in a fold with the given genotype. A balanced number of windows are drawn from the first half of each individual’s recording. From table 3, genotype-folds have 3, 4, or 5 individuals. For genotype-folds with 3 individuals, each individual contributes 13,333 or more windows, for genotype-folds with 5 individuals this is 8,000 windows.

The dictionary is initialized randomly, as in k-means, where the centroid waveforms are initialized as the first *P* points in *K* uniformly selected windows, chosen without replacement. A max of 300 iterations is chosen, but the dictionaries meet the other stopping criterion first, which is the mean squared difference after the centroid update is less than or equal to a tolerance value: 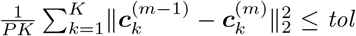. The tolerance *tol* is set to 10^−4^ times the pointwise variance of the training windows.

#### 3.1.4 Bag-of-Waves Representation

The bag-of-waves features are created by performing the assignment and collection steps to windows taken from a longer segment, converting the assignments into simple waveform counts. A segment 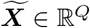 is broken into *L*-length windows. They could be non-overlapping for simplicity, or overlap with a stride of *L* − *P* + 1 to ensure matching of all possible shifts. For simplicity we consider non-overlapping and assume the segment length is truncated to a multiple of the window length, such that *M* = *Q/L* ∈ ℕ is the number of windows and 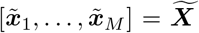. Applying the assignment and collection steps, equation 9 and equation 10, yields the counts *z*_*k*_ = |ℐ_*k*_|, *k* ∈ {1, …, *K*} forming the bag-of-waves vector ***z*** = [*z*_1_, …, *z*_*K*_] ∈ {0, …, *M*}^*K*^. For multiple dictionaries 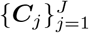, the bag-of-waves vector for each dictionary are concatenated ***z*** = [***z***^(1)^, …, ***z***^(*J*)^] ∈ {0, …, *M*}^*D*^, where ***z***^(*j*)^ is the bag-of-waves for the *j*th dictionary and 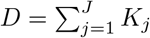 is the dimension of the bag-of-waves total number of waveforms.

##### Specifics for our dataset

We consider 1 hour segments resulting in *Q* = 921, 600 given the 256 Hz sampling rate. With *L* = 512 (2 s windows), this is *M* = 1, 800 windows per segment. For each of the *J* = 6 genotypes, 480 one-hour segments are selected from individuals with a given genotype and fold, with an equal number of segments per individual. Segments are drawn uniformly (possibly overlapping) from the second half of each individual’s recording, which ensures that if an individual was in the dictionary training fold then unseen windows are used for constructing the bag-of-waves vectors. For genotype-folds with 3 individuals, each individual contributes 160 one-hour segments, for genotype-fold with 4 individuals this is 120 one-hour segments, and for genotype-folds with 5 individuals this is 96 one-hour segments. As some individuals have recordings of shorter duration these segments overlap.

Additionally, to test whether classifiers trained using one-hour segments generalize to shorter time windows we also extract 2880 10-minute segments (each containing 300 windows) for each of the *J* = 6 genotypes. This amounts to the same duration per individual, taken also from the second half the recordings.

#### 3.1.5 Bag-of-Spectra Baseline

To assess whether the time-domain representation of the waveform is useful, we consider a baseline that assigns windows to clusters based on their spectra. That is for each window we compute a power spectral density as the squared magnitude of the discrete Fourier transform via the Fast Fourier transform (FFT). These are then normalized to sum to one (a discrete probability mass function across the sampled frequencies) and the square-root is taken. The processing is equivalent to simply using *ℓ*_2_-normalized FFT magnitude vectors. K-means is applied to cluster them. The spectra of new windows are processed in the same way and assigned based on the nearest centroid, with counts of centroid assignments forming a bag-of-spectra (BOS). The BOS uses the same data and hyper-parameters as BOW (except there is no use of *P*). Subsequent modeling is equivalent for both BOS and BOW.

### 3.2 Classification Models

We now consider modeling the relationship between bag-of-waves phenotypes and the genotype. For training, each individual *s* ∈ 𝒮_train_ has a genotype label *y*_*s*_ ∈ 𝒴 and a set of bag-of-wave segments. The training set label-feature pairs becomes 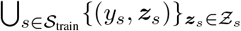, and the bag-of-waves alone is 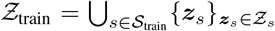 with.

#### 3.2.1 Feature Transformation via TFIDF

Before training a classifier, TFIDF weighting [33] and *ℓ*_2_-normalization is applied to the bag-of-waves. Specifically, the weighting factor *w*_*k*_ for *k* ∈ {1, …, *D*} is estimated from the training set as

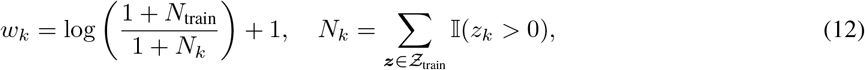

where *N*_*k*_ is the number of segments that the *k*th waveform is nonzero and 𝕀(·) is the 0-1 indicator function that is 1 if the logical argument is true. A feature vector after the TFIDF transformation is then

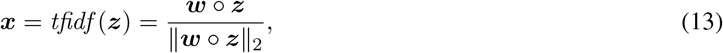

where ∘ indicates the element-wise multiplication and ***w*** = [*w*_1_, …, *w*_*D*_]. Applying this function to the bag-of-waves for individual *s* yields 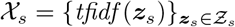. Note that this transformation is unsupervised.

#### 3.2.2 Logistic Regression

We consider logistic regressions models, which provide linear classifiers. For the case of binary classification |𝒴| = 2, assuming without loss of generality that the labels are 𝒴 = {0, 1} and *y* = 1 is the class of interest, the logistic regression model is

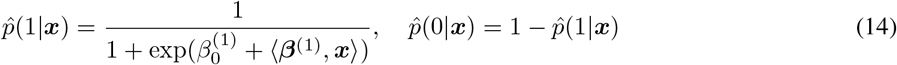

where 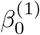 is the bias, 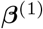 is the vector of coefficients. For |𝒴| *>* 2, the classifier is a multinomial logistic regression model

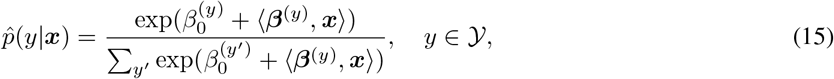

with biases 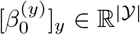 and coefficients [***β***^(*y*)^]_*y*_ ∈ ℝ^|𝒴|×*D*^. In either case, the parameters are optimized to maximize the log-likelihood of the model,

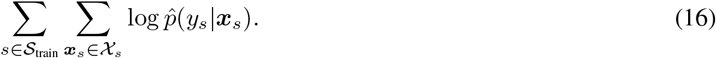

The multinomial model is overparametrized (only *C* − 1 bias and vectors of coefficients are needed). To ensure a unique solution, we penalize the magnitude of coefficients, via 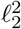 regularization, as in ridge regression [47], which is also necessary for correlated feature dimensions. Correlated features are possible if similar waveforms are in different class-conditional dictionaries as they will be selected as best matches for the same windows. The optimization of the parameters with the regularization is cast as the minimization of the following cost function

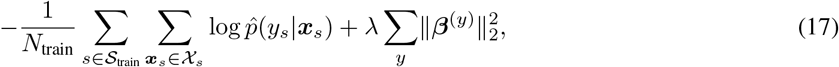

where 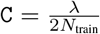 is the hyper-parameter and the summation in the second term is taken over all coefficient vectors.

#### 3.2.3 Pooled Counts

For inference on an individual *s* in either the validation and testing sets, the bag-of-waves occurrence count vectors are pooled across all segments and transformed

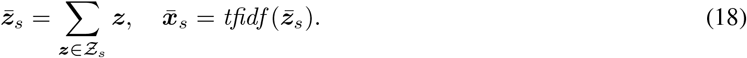

After this transformation, the validation set is denoted 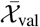, and the test set is denoted 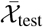. The model’s probability of a class *y* is 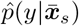.

Additionally, unpooled bag-of-waves features computed from different segments lengths can be used to test the segment-wise classification. The count across a segment of length *T* can be converted to occurrence rates, but due to the normalization in equation 13, 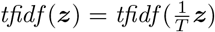, and the scaling does not change the feature. Thus, we will refer to waveform occurrence rates rather than counts.

#### 3.2.4 Factorized Classification

In general, predicting the joint genotype of an individual may be difficult as the number of genes of interest increase, and each gene of interest may have multiple alleles as classes. Likewise, a broad genotype associated to strain or genetic background can have multiple strains as classes. Mathematically, the joint genotype ***y*** = [*y*_1_, …, *y*_*G*_] consists of *G* component genotypes *y*_*g*_ ∈ 𝒴_*g*_, where |𝒴 _*g*_| is number of classes for the *g*th component. While each of the component genotypes *y*_*g*_ could be predicted independently 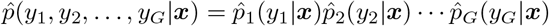, we consider factorized prediction where some genotypes are conditionally predicted based on other independently predicted genotypes. Specifically, for two components (*G* = 2) with the second component conditional on the first the model is

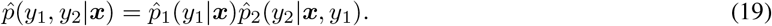

##### Specifics for our dataset

In this work *G* = 2 and we specifically focus on the case that *y*_1_ is the background strain with |𝒴_1_| = 3 classes and *y*_2_ is the TSC-genotype |𝒴_2_ | = 2. We train models for the joint 6 class problem, two independent models: one that predicts background strain and one for TSC-genotype, and TSC-genotype prediction conditional on the background strain.

#### 3.2.5 Nested Cross-validation for Hyper-parameter Selection and Leave-one-out Model Training

Here, we use an overall two-fold division of individuals for feature extraction (dictionary training). Let 𝒮 = {1, …, *S*} denote the index set of individuals, then 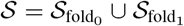, where 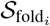 is the *i*th fold.

When one of the folds 𝒮_dict_ is used for feature extraction (training the waveform dictionaries per genotype), the other fold is divided into another training set portion 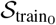 and a test set 𝒮_test_, such that 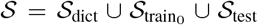. For a limited dataset, the genotype classification can be performed using leave-one-out (LOO) cross-validation. That is, a sequence of test sets consisting of one individual are formed 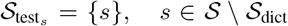, with the remaining individuals 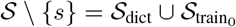 used for training and model selection.

With LOO performance evaluation, nested cross-validation is used for model selection: models are trained on 𝒮 _dict_ and a portion of 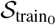 and validating on a held-out portion of 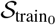. The motivation for only using 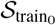 is that the feature distribution may differ between 𝒮_dict_ and 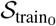, since the former was involved in the feature learning.

Stratified sampling is used to maximize the uniformity of the class distribution across the cross-validation folds, as implemented by `StratifiedGroupKFold` [48], where ‘Group’ refers segments of an individual. For *K*_CV_ internal folds, 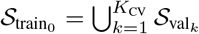. Then the individuals in the cross-validation training set for the *k*th fold is 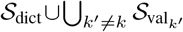 and the held-out validation fold is 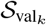.

To select the regularization hyper-parameter `C` a set of models is formed for each hyper-parameter value and cross-validation fold, and the hyper-parameter is selected that has the highest average classification accuracy (pooled with prediction per individual) across the validation folds, and if tied the highest average segment-wise accuracy. Then, a model is fit with this hyper-parameter model with the LOO training set, which is 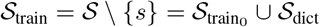, and tested on *s*. This is repeated for each individual in 𝒮 \ 𝒮 _train_, and then roles of the two-folds are reversed such that 𝒮_dict_ is the other fold.

##### Specifics for our dataset

For this dataset, we choose *K*_CV_ = 3 and select C from a grid of 15 logarithmically spaced vales from 10^−1^ to 10^4^, inclusive. In summary, dictionaries are trained from half of the individuals, models are estimated based on all but one individual and tested on the held-out individual. The models use one of two sets of features obtained by the dictionary learning. Finally, to check consistency of the entire genotype classification performance, five two-fold splits are created, but the feature analysis will focus on the dictionaries from the first split.

### 3.3 Feature Interpretation

The bag-of-waves representation is interpretable by design. The feature value is based on the occurrence rate of a waveform, and the waveforms corresponding to the classifier’s most important features can be visualized. Additionally, the spectra of important waveforms can be analyzed to identify key rhythms in the EEG records. We apply Shapley values, under an independence assumption [30, 31], as a computationally simple approach to compute instance-wise importance, and summarize the feature importance across all classification models using the same set of dictionaries by correlating the Shapley value and a class indicator. We propose to filter to those waveforms that have positive coefficients and whose counts also positively correlate with the class. Finally, we compute summary statistics about the waveform count and its spectrum.

#### 3.3.1 Shapley Values

In multinomial logistic regression, the log-odds for class *y* is

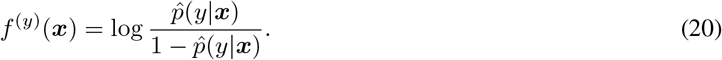

Assuming independent features [30, 31], the Shapley value for the *k*th feature is simply the product of the coefficient and the feature values difference from its mean

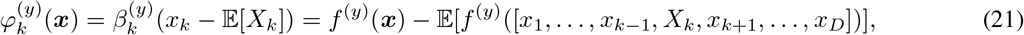

where *X*_*k*_ is the random variable corresponding to the *k*th feature. In practice, the expectation is estimated from the training set, 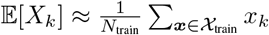. Shapley values are additive such that the prediction can be expressed as

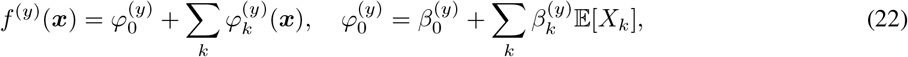

where 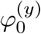 is the average prediction value for class *y*.

While Shapley values across the dataset can be summarized in multiple ways, such as their average magnitude, we propose the average class-signed Shapley value (ACSSV)

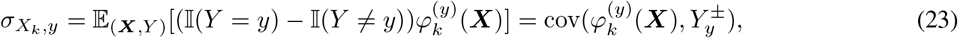

which is the covariance between the Shapley value 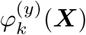 and the signed class indicator 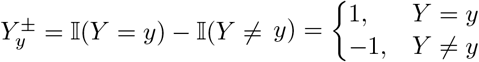. The covariance follows from the fact that 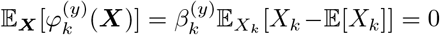, combined with the fact that the covariance is simply the expected product of two random variables when either of the random variables is zero mean.

As a covariance, the ACSSV will be high and positive when 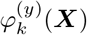 positively correlates with *Y* = *y*. This happens in two cases: first, when the coefficient 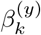 is positive and *X*_*k*_ positively correlates with *Y* = *y*, i.e., *X*_*k*_|*Y* = *y* is above the feature’s mean value and *X*_*k*_|*Y* ≠ *y* is below the feature’s mean, and second, when the coefficient is negative and the feature negatively correlates *Y* = *y*. Conversely, the ACSSV will be high and negative when either the coefficient is positive and the feature value in class *y* is below the feature’s mean value, or when the coefficient is negative and the feature value is above the mean. Thus, for the multinomial logistic regression model, the Shapley value feature analysis needs to consider both the sign of the coefficient and the sign of the ACSSV, as either alone is in insufficient to interpret a feature’s usage by a model.

In the case of cross-validation performance, different individuals have different classifiers, and for the LOO case, a different classifier for each individual. Nonetheless, all individuals in the same fold share the same dictionary, and the goal is to understand which of the bag-of-waves features, corresponding to waveform, is most important. Additionally, to compute the Shapley values we consider the pooled feature representations after applying the TFIDF transformation to the pooled bag-of-waves count vectors. Note that the TFIDF transformation is dependent on the training set statistics. Let 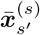 and 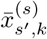 denote the feature vector and the *k*th feature value for *s*^′^ when trained with *s* held out.

When fold-*i, i* ∈ {0, 1} is used to form the dictionary, 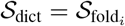, then individuals in 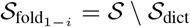 are tested individually. Let ***β***^(*y,s*)^ denote the coefficients for the *y*th class for a model trained with *s* held out (since *s* is specific to the fold, *i* is does not need to be specified). With this notation, the average class-signed Shapley value of the *k*th feature in the *i*th fold for class *y* is

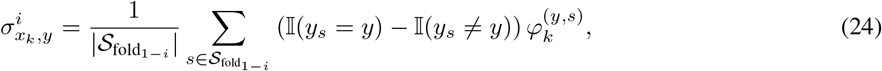

given the Shapley values

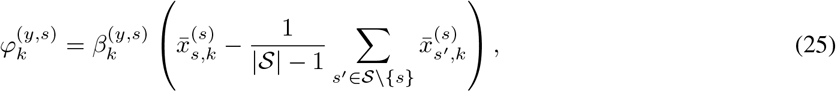

where 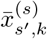 denotes the *k*th feature value for *s*^′^ when trained with *s* held out.

For the factorized classifier, used on our dataset, ACSSV is computed for *y*_2_ (the TSC-genotype) given *y*_1_ (the background strain) as

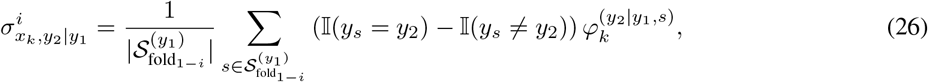

where 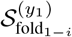 denotes the subset of the fold from class *y*_1_ and the Shapley values are

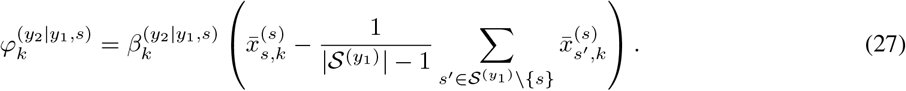

#### 3.3.2 Waveform Feature Analysis

The power spectral density function is estimated using the Welch periodogram method, averaging the squared magnitude of the discrete Fourier transform taken from overlapping windows. Starting from the 1 second waveforms, which is 256 samples, we extract two windows of length 243 (95%) extracted with 90% overlap (230 samples), which corresponds to a stride of 13. The default Hann function is applied, and each is zero-padded to 1024. This creates relatively smooth spectra for the relatively short duration waveforms.

The occurrence rates of waveforms (in units of counts per minute) for each class, outside of specific classes, and overall is computed across the training set.

## 4 Results

We present quantitative performance results and feature analysis of the proposed methodology for the background strain-TSC genotype EEG mice dataset (*n* = 45).

### 4.1 Classification Results

Genotype classification results are reported for background strain, TSC-genotype, TSC-genotype given background strain, and finally joint genotype.

#### 4.1.1 Background Strain Prediction

We first examine the accuracy of the BOW-based background-strain classifier in terms of the confusion matrix in Figure 2 computed for all *n* = 45 individual mice, using the original split. The accuracy per strain are 71%, 71%, and 79% for BXD87, DBA2, and C57B6, respectively.

**Figure 2:**
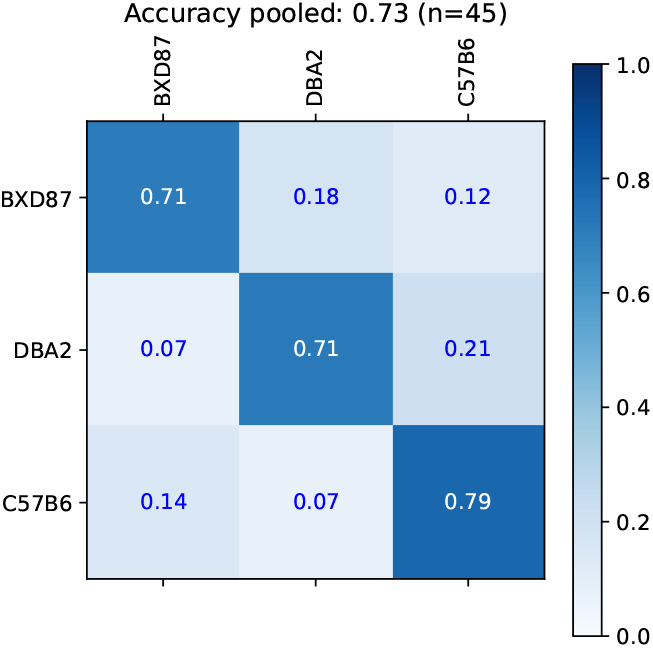
Confusion matrix for the prediction of background strain using BOW. The rows are the true class and columns are predicted class. The diagonal entries are the accuracy per strain of 71%, 71%, and 79% for BXD87, DBA2, and C57B6, respectively.

The mean and standard deviation of accuracies across the 5 two-fold splits are shown in Table 4, along with the bag-of-spectra (BOS) baseline. BOW and BOS achieve an average of 70% and 69%, respectively. This is well above 38% (17/45) achievable by the naive predictor that always selects the majority BXD87 strain. BOW and BOS perform comparably for this task, indicating that the distribution of spectra can distinguish individuals from different strains.

**Table 4:** Background Strain Classification Accuracy (3 classes).

| Method | BXD87<br>n=17 | DBA2<br>n=14 | C57B6<br>n=14 | Average<br>n=45 |
| --- | --- | --- | --- | --- |
| BOW | 0.63±0.05 | 0.73±0.08 | 0.77±0.04 | 0.70±0.02 |
| BOS | 0.65±0.04 | 0.65±0.03 | 0.77±0.04 | 0.69±0.01 |

Taking predictions for one-hour segments with BOW classifier without pooling yields a significantly lower average accuracy of 0.63 ± 0.01. Using only 10-minute segments the BOW classifier achieves 0.53 ± 0.01, which, while lower than longer windows, is well above chance. A classifier trained specifically for shorter windows may yield higher performance.

#### 4.1.2 TSC-Genotype Prediction

As shown in Figure 3, cla sification models that directly model TSC-genotype had chance or worse level performance, indicating that the two TSC-genotypes are not linearly separable in the feature spaces when ignoring the background strain.

**Figure 3:**
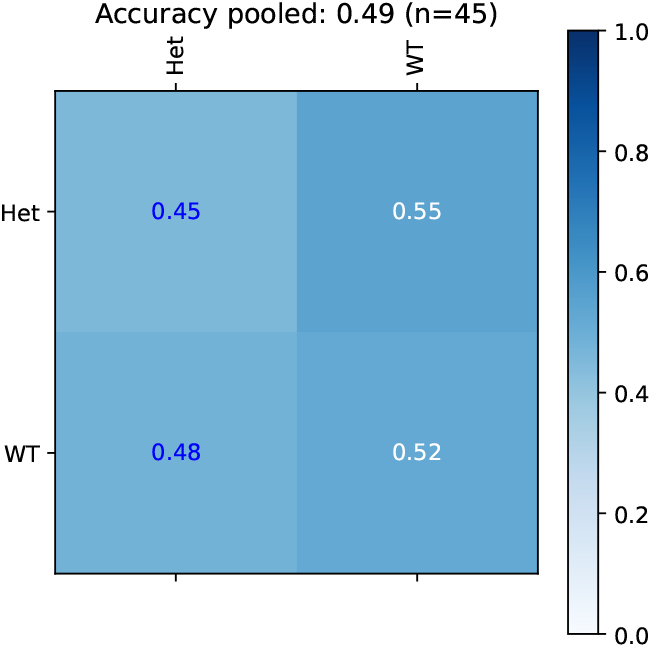
TSC-genotype prediction is at a chance level performance.

Given the true background-strain class, we train a class-conditional classifier to predict the presence of the *TSC1* knockout in heterozygous individuals (Het) versus wild-type (WT) individuals. The confusion matrix for the BOW classifier is shown in Figure 4 for the original split. The TSC-genotype classification are reasonably accurate for the DBA2 and C57B6 strains, achieving a sensitivity of 67% and specificity of 100% for DBA2-Het, and a sensitivity of 83% and specificity of 100% for C57B6-Het. However, for the BXD87 strain, the TSC-genotype classification is at chance level.

**Figure 4:**
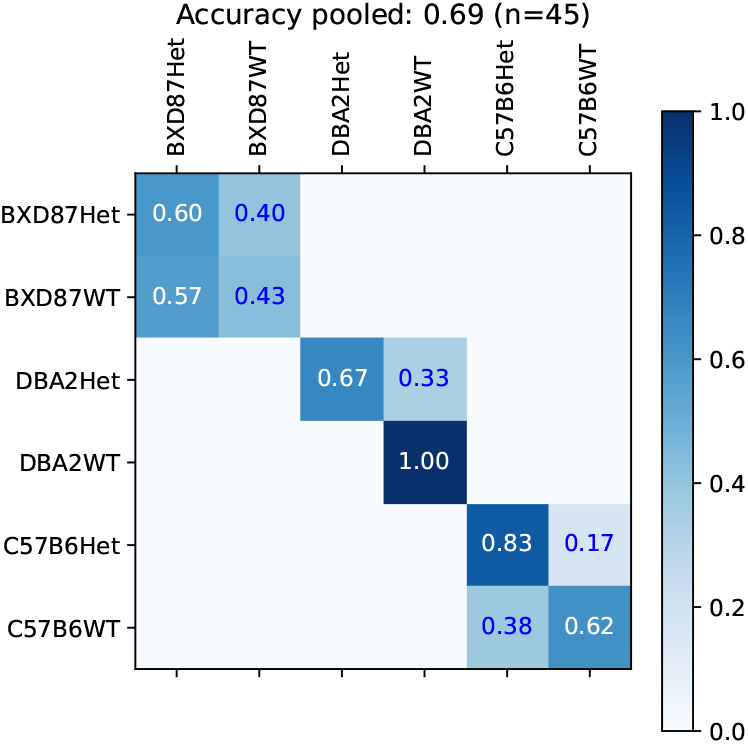
TSC-genotype prediction given background strain.

The average and standard deviation of accuracy across the 5 two-fold splits are in Table 5. The BOW classifiers show moderate prediction accuracy of TSC-genotype 86% and 67% for DBA2 and C57B6, respectively, but below chance accuracy for BXD87. The naive classifier performance is 59% (10/17), 57% (8/14), and 57% (8/14), for BXD87, DBA2, and C57B6, respectively. In comparison, the BOS classifier has lower performance (beyond 1 standard deviation) with accuracies of 79% and 53% for DBA2 and C57B6, respectively.

**Table 5:** TSC Genotype Classification Accuracy (2 classes).

| Method | BXD87<br>n=17 | DBA2<br>n=14 | C57B6<br>n=14 | Average<br>n=45 |
| --- | --- | --- | --- | --- |
| BOW | 0.46±0.05 | 0.86±0.05 | 0.67±0.04 | 0.65±0.03 |
| BOS | 0.53±0.04 | 0.79±0.00 | 0.53±0.10 | 0.61±0.03 |
| Chance | 0.59 | 0.57 | 0.57 | 0.58 |

The accuracy is computed using predictions from the logistic regression models based on a probability threshold of 0.5. To achieve a desired specificity or sensitivity the threshold could be adjusted, and the receiver operating characteristic (ROC) curve traces the sensitivity and specificity across a range of thresholds. Using the classifiers predicted probabilities for the TSC-genotype we compute ROC curves for each of the 5 splits. These curves, and the area under the curve (AUC), are reported in Figure 5 along with with the average ROC based on merging [49, 50]. The curves indicate that the BOW classifiers are superior to BOS classifiers across the range of sensitivities, and detecting the TSC-genotype is more difficult for the C57B6 strain.

**Figure 5:**
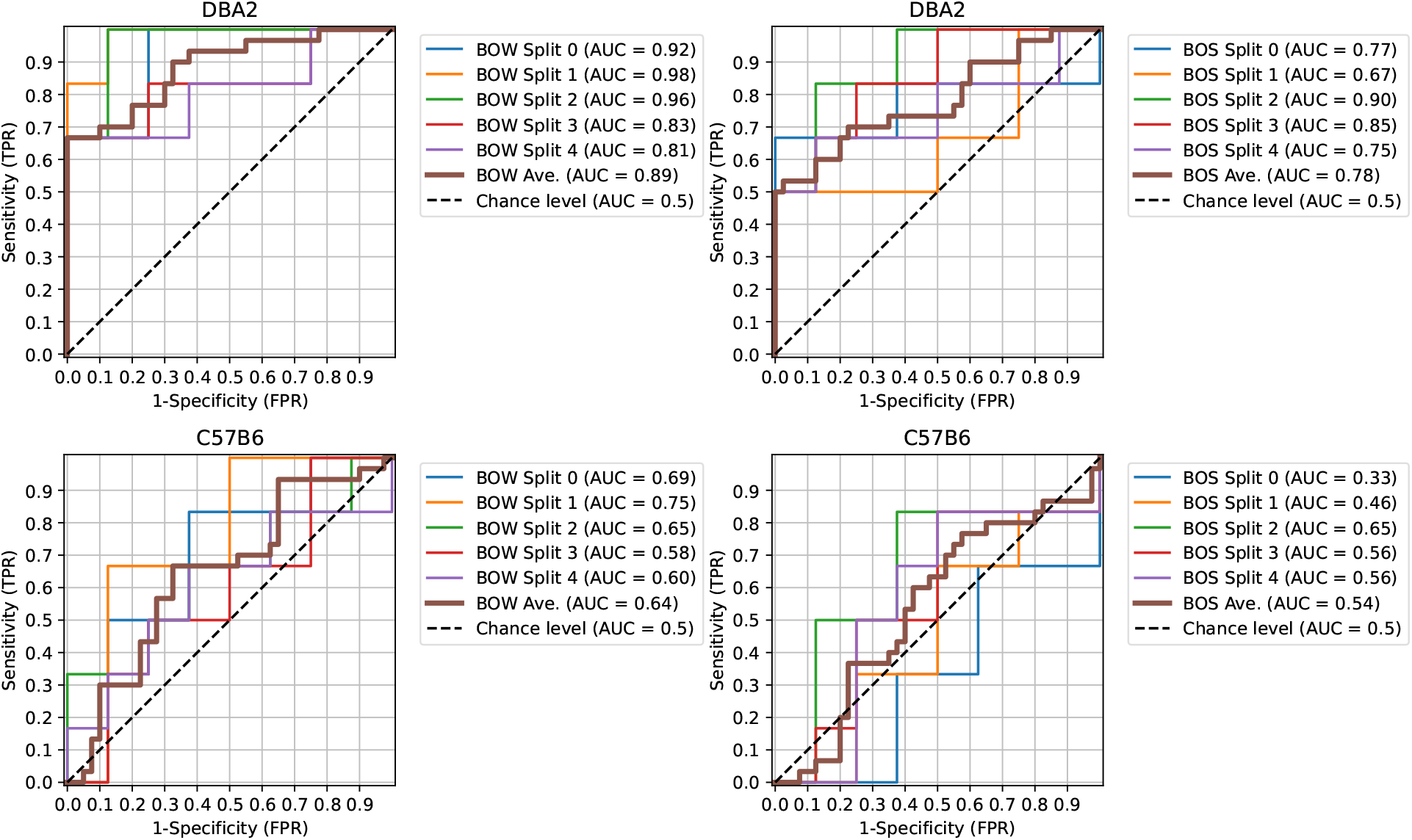
Receiver operating characteristic (ROC) curve for TSC-genotype prediction given background strain. (Left) BOW classifiers. (Right) BOW classifiers.

The sensitivity and specificity for the strain-specific BOW TSC-genotype classifiers for the pooled, hourly, and 10- minute predictions are in Table 6. When using the pooled representation, the sensitivity in detecting *TSC1* knockout (Het) is 67% for both background strains. For the DBA2 strain, WT is easier to identify yielding higher specificity. This indicates that longer-term recordings are generally more sensitive and specific, but one-hour recordings are reasonably accurate. Using the hourly representation, only the sensitivity for the DBA2 strain is maintained but the other performance measures drop. Using only a 10 minute segment, the specificity drops in both cases, but the sensitivity is actually higher for DBA2 indicating a bias towards Het. This bias is not unexpected due to differences in the count-based features for shorter windows.

**Table 6:** TSC Genotype BOW Classifier’s Sensitivity and Specificity.

|  | DBA2 |  | C57B6 |  |
| --- | --- | --- | --- | --- |
|  | Het<br>Sensitivity<br>n=6 | WT<br>Specificity<br>n=8 | Het<br>Sensitivity<br>n=6 | WT<br>Specificity<br>n=8 |
| Pooled | 0.67±0.12 | 1.00±0.00 | 0.67±0.12 | 0.67±0.07 |
| Hourly | 0.68±0.02 | 0.92±0.03 | 0.57±0.09 | 0.62±0.03 |
| 10 min. | 0.76±0.06 | 0.67±0.03 | 0.48±0.08 | 0.60±0.04 |

#### 4.1.3 Joint Genotype Prediction

Jointly predicting one of the six strain-TSC genotypes is the most challenging task. We compare the proposed factorized classification (FC) approach, where the prediction is made by combining the strain classifier with the strain-conditional TSC-genotype classifier, to the joint classifier. The confusion matrices, on the original split, for the joint and FC classifier using the BOW representation are shown in Figure 6. The accuracies for joint and factorized classification are 36% and 47%, respectively, both are well above the chance given by a naive predictor with accuracy of 22% (10/45). We examine whether the joint genotype classifications reliably identify the Het TSC-genotype. Considering the factorized classifier, for DBA2, the sensitivity is 66% and the specificity is 100%, which is the same classification given the true background. For C57B6, the sensitivity is 67%, but the specificity is 38%, as some C57B6WT individuals are classified as BXD87Het. For the joint classifier, for DBA2, the sensitivity is 67% and the specificity is 88%, as some DBA2WT individuals are classified as BXD87Het. For C57B6, the sensitivity is 34% and the specificity is 67%.

**Figure 6:**
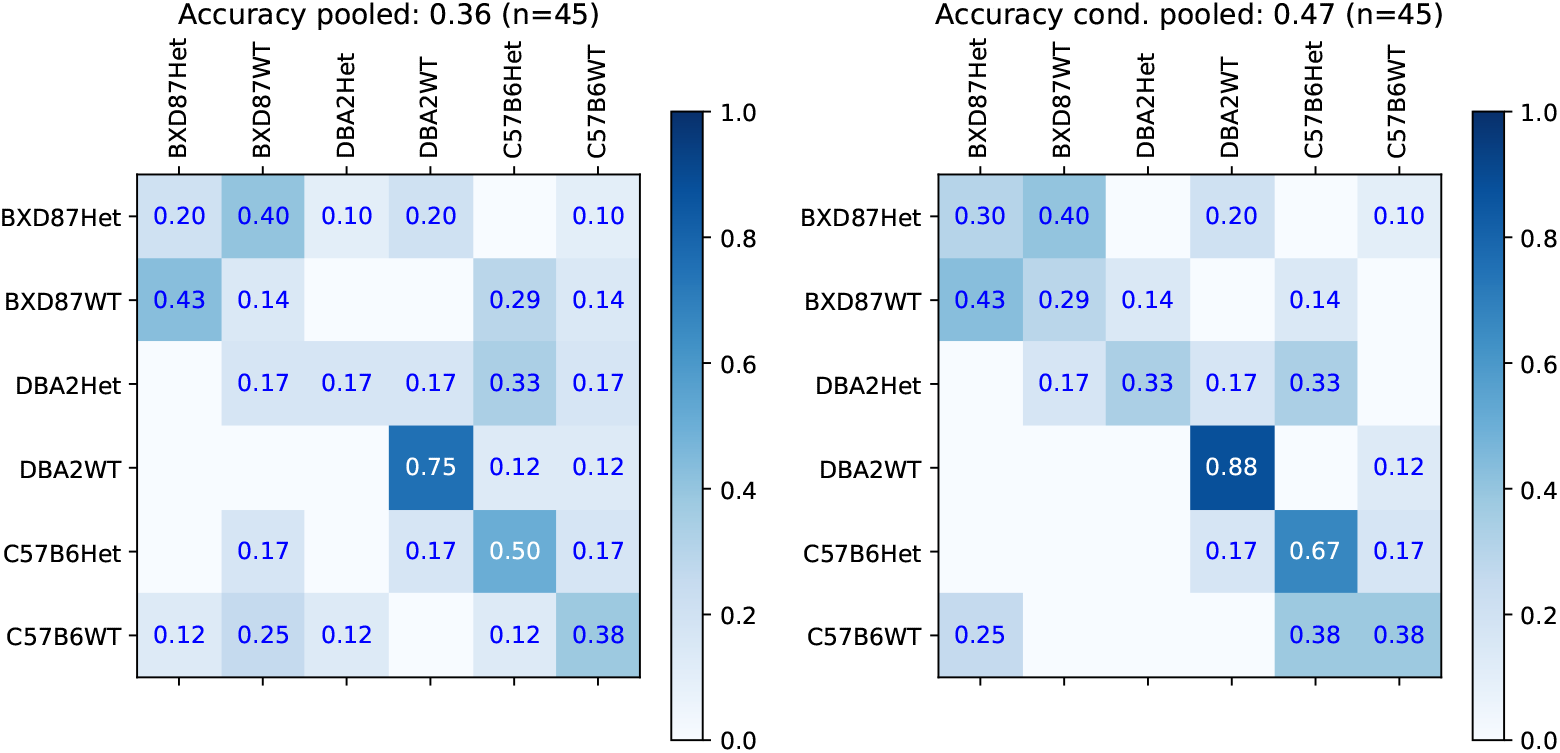
Joint genotype prediction across 6 types using either a single classifier (left) or the factorized classification using strain classifier combined with the TSC-genotype prediction given strain classifier (right).

The mean and standard deviation across the 5 splits for both BOW and BOS representations and joint and FC classification is shown in Table 7. The factorized models (FC) are superior to the joint models, and BOW and BOS have comparable performance. This indicates that the relationship between the features and the TSC genotype is not linear. Within a strain, a linear model can better separate the TSC genotypes (Het and WT classes).

**Table 7:** Genotype Classification Accuracy (6 classes).

| Method | Joint | FC |
| --- | --- | --- |
| Chance | 0.22 | 0.22 |
| BOS | 0.35±0.03 | 0.45±0.04 |
| BOW | 0.35±0.03 | 0.43±0.04 |

### 4.2 Waveform Feature Analysis

To interpret the genotype classification models we identify the waveforms associated to the highest average class-signed Shapley values (ACSSV) with positive average coefficients. The restriction to positive average coefficients is because a waveform occurrence feature may have a high average class-signed Shapley values but be negatively associated with a class. The ACSSV are interpreted by examining each waveform’s occurrence rate within a class and outside of the class, along with its average coefficient. ACSSV can be large and positive when either the waveform occurs more often in a class and the coefficient is positive, or if the waveform occurs less often in a class and the coefficient is negative.

#### 4.2.1 TSC Genotype Classifiers

Figure 7 illustrates the relationship between the ACSSV, average coefficient, and difference of waveform occurrence rates for the TSC genotype classifier for DBA2 and C57B6 strains across both folds. We visualize the waveforms, their spectra, and report their spectral peaks and occurrence rates in Figure 8 for DBA2 and in Figure 9 for C57B6. For DBA2, in Fold 0, the top-3 Het waveforms are transient waveforms that occur more often in Het than WT. The 4th and 5th Het waveforms are irregular rhythms that are from the DBA2Het dictionary (ID begins with 3), with similar occurrence rates in both classes. The WT waveforms are rhythmic signals with peak frequencies 6.2–8 Hz. In Fold 1, none of the Het waveforms appear more often in Het than WT, but all the WT waveforms appear more often in WT than Het and are rhythmic waveforms with peaks frequencies 5.5–7.8 Hz.

**Figure 7:**
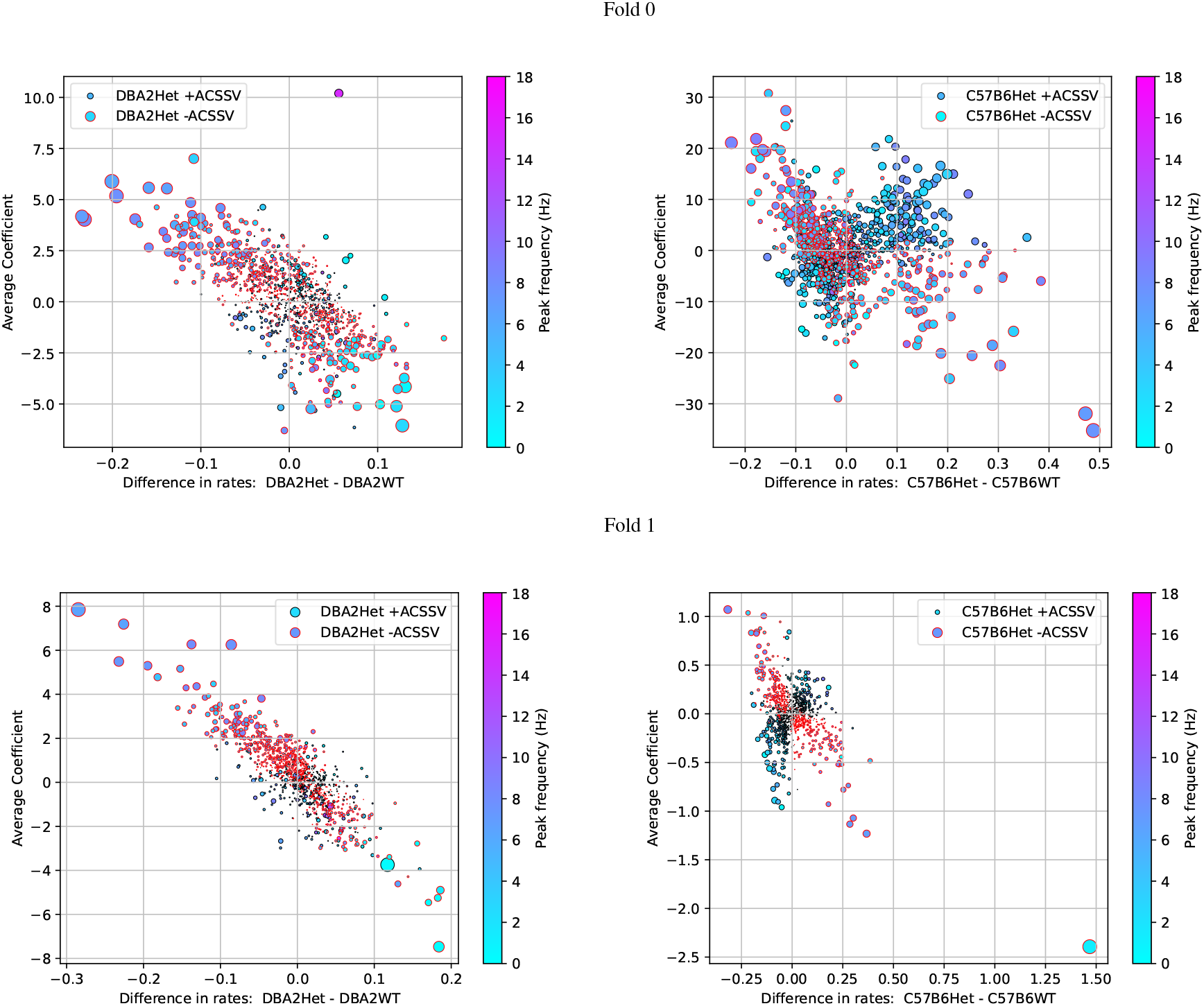
Scatter plots showing the relationship between the average class-signed Shapley values (ACSSV), average coefficient, and difference of waveform occurrence rates (count per minute) for the TSC genotype classifier for DBA2 and C57B6 strains across both folds. Marker size is proportional to the magnitude of the ACSSV (relative to maximum in a given plot), and black outlined points correspond to waveforms whose Shapley values positively correlate with the TSC genotype, whereas red outlines negatively correlate. The marker color corresponds to the peak frequency of the waveform. (Left) For DBA2 there are relatively few waveforms that have increased occurrence in DBA2Het and positive coefficients indicative of the Het genotype, which means the classifiers instead rely on the reduced rate of occurrences of waveforms indicative of the wild type for classification. (Right) For C57B6 there are more waveforms with increased rate for C57B6Het and positive coefficients indicative of the Het genotype.

**Figure 8:**
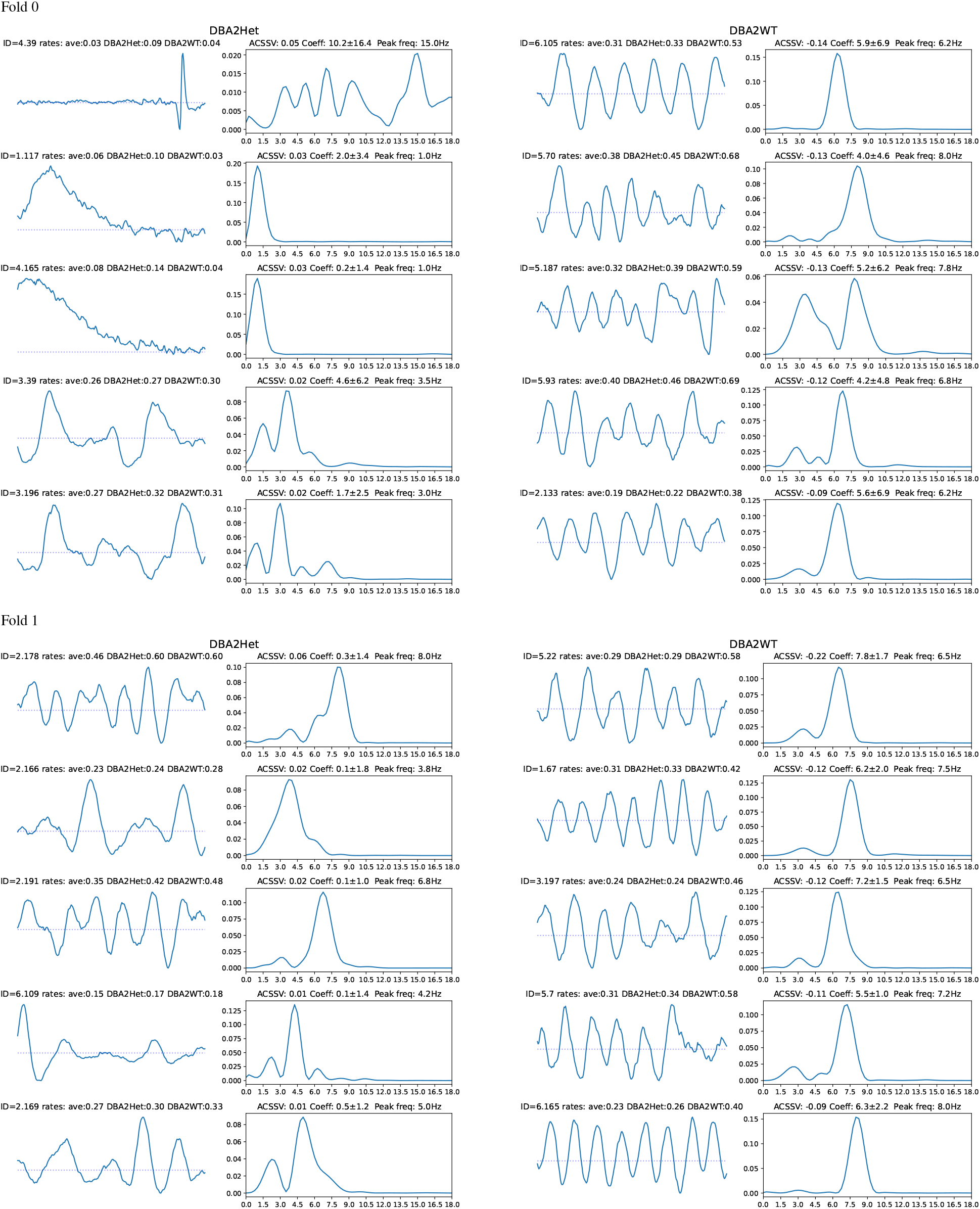
Bag-of-waves features analysis with Shapley values for the TSC-genotype classifier given DBA2 strain. The waveforms (1 s duration) and their spectra with the top-5 average class-signed Shapley values for Het (Left) and WT (Right) with positive coefficients.

**Figure 9:**
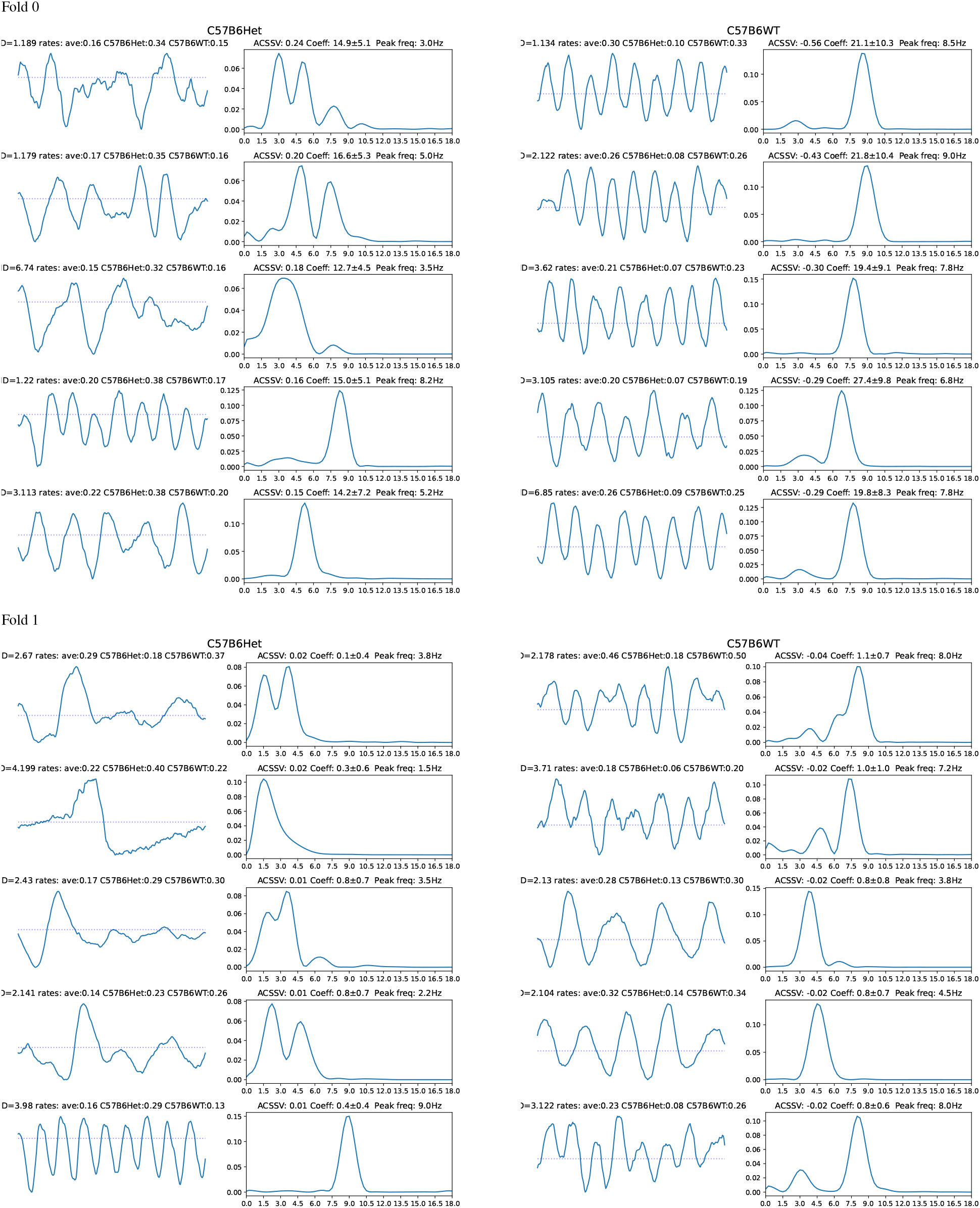
Bag-of-waves feature analysis with Shapley values for the TSC-genotype classifier given C57B6 strain. The waveforms (1 s duration) and their spectra with the top-5 average class-signed Shapley values for Het (Left) and WT (Right) with positive coefficients.

For C57B6, in Fold 0, the top-3 Het waveforms are irregular rhythms and the 4th and 5th are regular rhythms. The WT waveforms are all rhythmic signals with peak frequencies 6.8–9 Hz. In Fold 1, the first 4 Het waveforms are transients or irregular rhythms (only the 2nd appears more often in Het than WT) and the 5th waveform is a rhythm. All the WT waveforms appear are rhythmic waveforms (the 1st, 2nd, and 5th are slightly irregular) with spectral peaks frequency 3.8–8 Hz.

Across both strains, it can be observed that the waveforms indicative of the Het genotype (presence of *TSC1* knockout allele) are generally transients or irregular rhythms (occasionally rhythmic waveforms) that occur more often in Het than WT, whereas waveforms indicative of the wild type are generally rhythmic waveforms (sometimes involving bimodal spectra that are more irregular). Additionally, we note the higher ACSSVs and average coefficients for the waveforms in C57B6Het compared to DBA2Het, which indicates that for DBA2 the TSC-genotype classifiers are actually relying on waveforms indicative of wild type. If the Het TSC-genotype creates more diverse phenotypes, then this strategy is logical as the ‘normal’ wild-type waveforms may be stereotyped providing more reliable features.

Looking at the correspondence with the scatter plots in Figure 7, we note that the first waveform in Fold 0 for the DBA2Het is the high-frequency transient has a difference in rate of 0.05 and an average coefficient of 10.2, which appears as an outlier in the top-right quadrant. This transient occurs with a relatively low occurrence rate corresponding to an average interval of 11 minutes (0.09 times per minute) in DBA2Het mice but (0.09 times per minute) compared an interval of 25 minutes (0.04 times per minute) in DBA2WT. The next two waveforms are slow waves that peak at the high-pass cut-off frequency, they are also are relatively low occurrence rate. Finally, we note that the outlier point in Fold 1 for C57B6 in Figure 7 is not shown as it occurs more frequency in C57B6Het but the classifier assigns a negative coefficient to it, meaning it is indicative of C57B6WT.

Figure 10 shows the spectra of the top waveforms for the Het and WT classes given the strain. It is easier to see spectral shifts for the DBA2 strain, with the DBA2Het class corresponding to lower frequency waveforms and the DBA2WT waveforms peaking 3 Hz or 6–8.5 Hz. For C57B6, it is more difficult to see the consistent spectral shifts across both folds.

**Figure 10:**
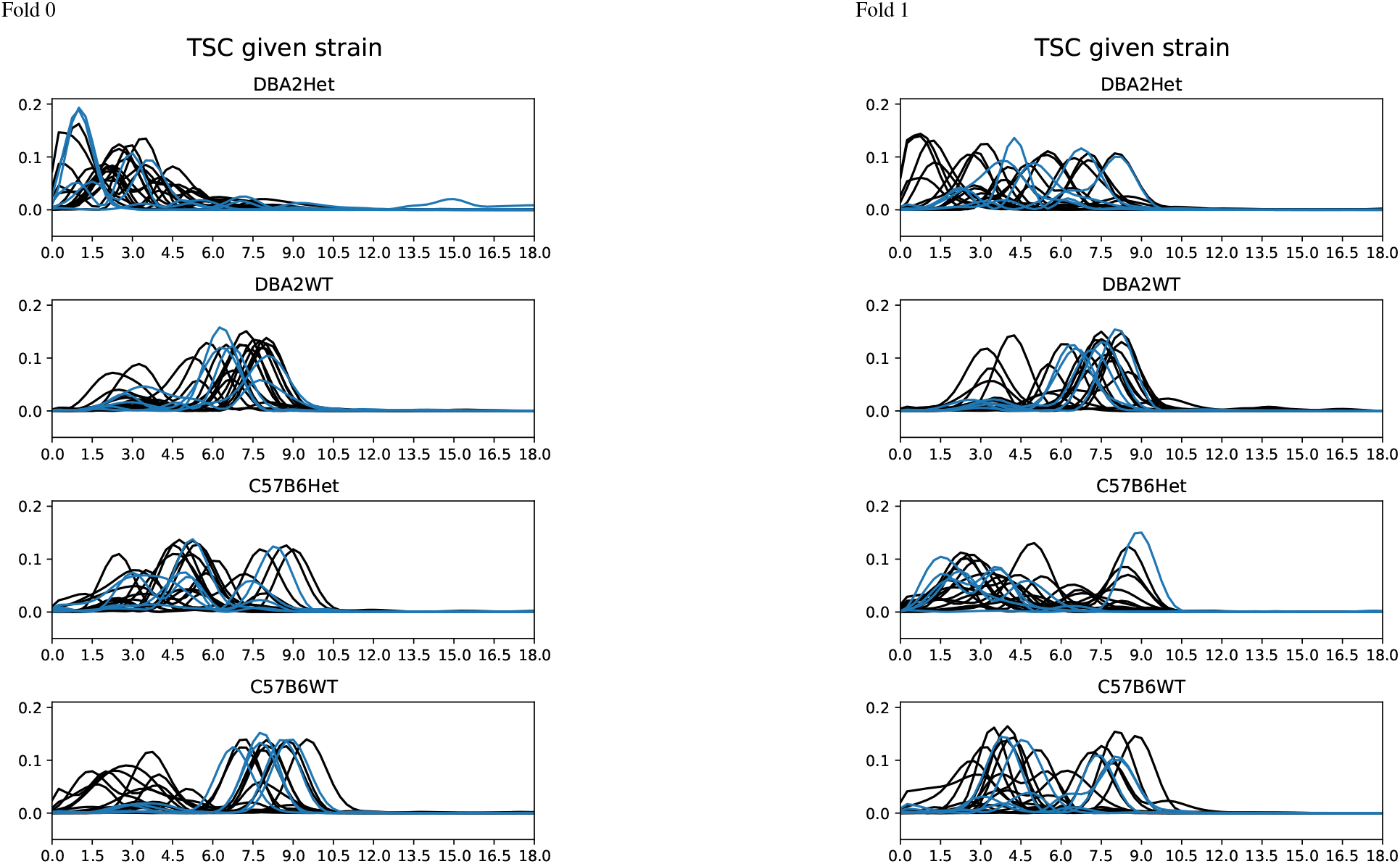
Spectra of waveforms with top-20 average class-signed Shapley value for strain-conditional TSC-genotype classifier, heterozygous (Het) versus wild-type (WT), for DBA2 and C57B6 strains. Top-5 waveform spectra are blue. For DBA2Het, waveforms tend to be lower frequency or peak at different frequencies than the 5–9 Hz in DBA2WT. In Fold 0, there is one with broad spectra (impulse-like transients in time-domain). The critical waveforms for WT have spectra for both strains that are remarkably similar. For C57B6Het, the peak frequencies peak higher in between the two clusters in WT. For C57B6Het, there are less waveforms with strong peaks between 6–7.5 Hz.

## 5 Discussion

Our results show that information from both inbred genetic background and TSC-genotype can be determined from bag-of-waves EEG biomarkers. Interestingly, the TSC-genotype could not be reliably classified with a linear model without knowledge or an estimate of the background strain, while in conditional models, the TSC-genotype could only be reliably determined in two of the strains (DBA2 and C57B6). The lack of predictive power in the BXD87 background is intriguing because this was the only strain background in which we observed spontaneous seizures, suggesting that neural activity in wild type mice was more similar to the knockouts, which we speculate is more disease like. These results highlight the important role genetic backgrounds play in determining phenotypes and disease. The classification rate for the background strain is 70%, which indicates that there are robust phenotypic markers in the EEG. The sensitivity and specificity for the *TSC1* knockout allele (Het) was relatively high, with sensitivities of 67% at a specificity of 100% for DBA2 and 67% for C57B6. The absence of any observed seizures (in video or through electrophysiological analysis) in mice with these genotypes, supports the non-ictal changes previously reported [10, 11]. Thus, despite lacking the overt spontaneous seizures of epilepsy, these mice have subtle phenotypes that can be detected automatically using the BOW approach.

These differences may also be manifested in behavioral activity in these freely behaving mice. Fundamentally, genetics drives the neurophysiological activity and behavior. Our analysis showed that pooling multiple 1 hour segments over at least 24 hours increases the accuracy, but hour long counts and even 10 minute counts still achieve above chance prediction. Future analysis could investigate the relationship between waveforms and behavior, whether waveforms most indicative are present during wake or sleep, locomotion, etc. In particular, the 2nd and 3rd waveforms associated with the *TSC1* knockout in DBA2 strain (DBA2Het) in Fold 0 (Figure 8) could possibly be artifacts or be associated with slow waves associated with wake/sleep cycles [51, 52] and observed in anesthetized mice [53].

### 5.1 Disease relevance of the novel mouse panel

In humans, single *TSC1* or *TSC2* mutations can have differential effects between affected individuals, and even **identical** inherited variants can result in different disease profiles. This indicates significant heterogeneity in patient outcomes, the mechanisms for which are currently unknown; having a biomarker for this difference in prognosis will allow us to make more accurate predictions about disease severity and clinical outcome in affected individuals. To this end, we use here a model system of this heterogeneity to improve our ability to probe the basic biology of TSC and explore the neural network differences underlying seizures. Our novel mice have a germline heterozygous *TSC1* knockout which, when introduced to controlled genetic background diversity, faithfully recapitulates patient heterogeneity in TSC results with the clinically relevant gene dosage and expression pattern. The ability to automatically detect subtle genotype-specific EEG phenotypes across diverse genetic backgrounds underscores the translational potential of this panel for preclinical biomarker discovery. By leveraging a scalable and interpretable machine learning framework, this approach mirrors the complexity of human neurological disorders, where genetic heterogeneity and subclinical manifestations often obscure diagnosis. The design of the panel, which incorporates multiple strains across wild-type and disease-associated genotypes, offers a powerful analog to the diversity of the human population, allowing the identification of EEG characteristics that can be generalized between individuals or highlight genotype-specific vulnerabilities. This positions the BOW methodology not only as a tool for mechanistic insight but also as a bridge toward precision diagnostics in clinical neurology, particularly in cases where overt seizures are absent but underlying pathology persists or where genetic testing indicates the presence of a seizure-associated gene but prognosis is unclear.

### 5.2 Translational potential of BOW representations

While we proved the utility of the BOW approach on a unique animal dataset, the proposed machine learning approach and feature interpretation is general and can be applied to other EEG time-series classification problems. Applying the approach to human EEG data [54] to aid epilepsy diagnosis from recordings without any observed seizures is an obvious next step. In this case the dataset would consist of patients with epilepsy and controls. The records themselves may be shorter, or they may also be long-term recordings, but data sets from neurological clinics may have hundreds to thousands of individuals. A key difference is that human EEGs have many more channels, and waveform patterns associated to the condition may be spatially localized in some cases.

#### 5.2.1 Multichannel Generalizations

The bag-of-waves explored here considers only a single channel. In the multichannel case, it would be necessary to consider spatiotemporal waveform dictionaries [23]. The clustering approach, which assumes only a single prominent waveform in each window, may be inadequate for whole scalp models. Alternatively, a bag-of-waves may be computed on each channel separately using a shared (or channel-specific dictionary assuming standard EEG montage). Then a classifier could both the spatial pattern along with pattern across waveforms.

Another option would be to apply independent component analysis, filter to independent components (ICs) related to brain activity and discard artifact ICs classes [55], and compute bags-of-waves across all brain-related ICs. These could simply be pooled to compute a single representation. In this case, a fixed spatial pattern would not be available, but spatially organized pooling could be used to group ICs based on their spatial pattern across the scalp. Another option would be to cast the problem as multiple instance learning [56, 57], a paradigm for binary classification where the representation of each instance is an unordered set of points, at least one of which is positive for the positive class. Here the set of points would be the different EEG channels or independent components. For standard EEG montages, it would be possible to train channel specific dictionaries.

#### 5.2.2 Application to Other EEG Classification Tasks

Beyond genotype prediction, classification of long-term EEG is motivated in cases of diagnosis of other diseases, where an individual is assigned a single label. Additionally, the methodology is readily adapted to both shorter-term classification, where windows of EEG around known stimuli timings are classified, or segmentation, where each window is assigned to a class like in sleep staging, or seizure prediction. In particular, for *K* = 1 the shift-invariant k-means algorithm corresponds to a model for evoked potentials in EEGs that shift per trial [58]. Thus, the shift-invariant k-means provides a model with the flexibility to choose from *K* waveforms to approximate evoked potentials in response to stimuli.

### 5.3 Bag-of-Waves versus Bag-of-Spectra

We also produced a baseline approach that assigns the spectrum of each window to a set of centroids learned through k-means. The resulting bag-of-spectra performs comparably to bag-of-waves except on the TSC-genotype prediction given the strain, which is arguably the most interesting problem. The spectra representation, however, loses information about the phase that is needed to reconstruct the window, which makes the interpretation of the spectral patterns substantially more difficult, especially in the case of transient waveforms that are not rhythmic. Indeed, neurologists are trained to look at EEG in the time domain (not their spectra) precisely because waveform morphology is clinically meaningful. Thus, the bag-of-waves representation provides more natural interpretation.

## 6 Conclusion

We described and validated a machine learning approach to predict the genotype from long-term continuous EEG signals based on the occurrence rates of prototypical waveforms in EEG that are estimated via a shift-invariant clustering algorithm applied to segments of signals from each class. The bag-of-waves representation used here simplifies previous convolutional dictionary learning approaches, by simply finding one waveform optimally shifted and scaled for each window. We apply the proposed methodology to EEGs from mice where the classes correspond to genotypes of individual mice, of varying inbred strains and with or without the *TSC1* knockout associated with epilepsy. The genotypes of individuals are predicted from the occurrence of waveforms across multiple segments of single-channel EEG time series. We find reliable prediction of the mouse strain, and for two of the three strains, a strain-specific classifier can reliably determine the presence of the *TSC1* knockout. The waveforms most indicative of the genotypes are identified through Shapley values. The results show the promise and challenges in identifying EEG biomarkers for specific genotypes from single channel EEG. In particular, future work could adapt the methodology to larger datasets of human EEG [54], to aid the analysis of EEG for epilepsy diagnosis.

## 7 Acknowledgments

This research was supported in part through the use of Information Technologies (IT) resources at the University of Delaware, specifically the high-performance computing resources. Additional funding for this work came from NIH NINDS 5K22NS104230 and NIH NINDS 5R01NS134491 (awarded to AEH). We would like to thank Matthew Weston for his generous donation of the initial *TSC1*^tm1Djk^/J mice.

## References

[1] Paul L Nunez and Ramesh Srinivasan. Electric fields of the brain: the neurophysics of EEG. Oxford University Press, 2006.

[2] György Buzsáki. Rhythms of the Brain. Oxford University Press, 2006.

[3] Mina Amiri, Jean-Marc Lina, Francesca Pizzo, and Jean Gotman. High frequency oscillations and spikes: separating real hfos from false oscillations. Clinical Neurophysiology, 127(1): 187–196, 2016.

[4] Scott R Cole and Bradley Voytek. Brain oscillations and the importance of waveform shape. Trends in Cognitive Sciences, 21(2): 137–149, 2017.

[5] TH Bullock, MC McClune, and JT Enright. Are the electroencephalograms mainly rhythmic? assessment of periodicity in wide-band time series. Neuroscience, 121(1): 233–252, 2003.

[6] Jeffrey W Britton, Lauren C Frey, John L Hopp, et al. Eeg in the epilepsies. In Erik K St. Louis and Lauren C Frey, editors, Electroencephalography (EEG): An Introductory Text and Atlas of Normal and Abnormal Findings in Adults, Children, and Infants. American Epilepsy Society, Chicago, 2016.

[7] Matilde Leonardi and T Bedirhan Ustun. The global burden of epilepsy. Epilepsia, 43: 21–25, 2002.

[8] Peter B Crino, Katherine L Nathanson, and Elizabeth Petri Henske. The tuberous sclerosis complex. New England Journal of Medicine, 355(13): 1345–1356, 2006.

[9] Andrew R Tee, Brendan D Manning, Philippe P Roux, Lewis C Cantley, and John Blenis. Tuberous sclerosis complex gene products, tuberin and hamartin, control mtor signaling by acting as a gtpase-activating protein complex toward rheb. Current Biology, 13(15): 1259–1268, 2003.

[10] Davide Bassetti, Heiko J Luhmann, and Sergei Kirischuk. Effects of mutations in tsc genes on neurodevelopment and synaptic transmission. International Journal of Molecular Sciences, 22(14): 7273, 2021.

[11] Susanna MI Goorden, Geeske M Van Woerden, Louise Van Der Weerd, Jeremy P Cheadle, and Ype Elgersma. Cognitive deficits in Tsc1+/-mice in the absence of cerebral lesions and seizures. Annals of Neurology: Official Journal of the American Neurological Association and the Child Neurology Society, 62(6): 648–655, 2007.

[12] Michael S. Lewicki and Terrance J. Sejnowski. Coding time-varying signals using sparse, shift-invariant representations. Advances in Neural Information Processing Systems, pages 730–736, 1999.

[13] Evan Smith and Michael S Lewicki. Efficient coding of time-relative structure using spikes. Neural Computation, 17(1): 19–45, 2005.

[14] Thomas Blumensath and Mike Davies. Sparse and shift-invariant representations of music. IEEE Transactions on Audio, Speech, and Language Processing, 14(1): 50–57, 2005.

[15] Philippe Jost, Pierre Vandergheynst, Sylvain Lesage, and Rémi Gribonval. Learning redundant dictionaries with translation invariance property: the motif algorithm. In SPARS’05-Workshop on Signal Processing with Adaptive Sparse Structured Representations, pages 1–3, 2005.

[16] Chaitanya Ekanadham, Daniel Tranchina, and Eero P Simoncelli. Recovery of sparse translation-invariant signals with continuous basis pursuit. IEEE Transactions on Signal Processing, 59(10): 4735–4744, 2011.

[17] Boris Mailhé, Sylvain Lesage, Rémi Gribonval, Frédéric Bimbot, and Pierre Vandergheynst. Shift-invariant dictionary learning for sparse representations: Extending K-SVD. In 2008 16th European Signal Processing Conference, pages 1–5. IEEE, 2008.

[18] Roger Grosse, Rajat Raina, Helen Kwong, and Andrew Y Ng. Shift-invariant sparse coding for audio classification. In Proceedings of the Twenty-Third Conference on Uncertainty in Artificial Intelligence, pages 149–158, 2007.

[19] Austin J Brockmeier and Jose C Principe. Learning recurrent waveforms within eegs. IEEE Transactions on Biomedical Engineering, 63(1): 43–54, 2015.

[20] Mainak Jas, Tom Dupré la Tour, Umut Simsekli, and Alexandre Gramfort. Learning the morphology of brain signals using alpha-stable convolutional sparse coding. Advances in Neural Information Processing Systems, 30, 2017.

[21] Sebastian Hitziger, Maureen Clerc, Sandrine Saillet, Christian Bénar, and Théodore Papadopoulo. Adaptive waveform learning: a framework for modeling variability in neurophysiological signals. IEEE Transactions on Signal Processing, 65(16): 4324–4338, 2017.

[22] Quentin Barthélemy, Cédric Gouy-Pailler, Yoann Isaac, Antoine Souloumiac, Anthony Larue, and Jérôme I Mars. Multivariate temporal dictionary learning for EEG. Journal of Neuroscience Methods, 215(1): 19–28, 2013.

[23] Tom Dupré la Tour, Thomas Moreau, Mainak Jas, and Alexandre Gramfort. Multivariate convolutional sparse coding for electromagnetic brain signals. Advances in Neural Information Processing Systems, 31, 2018.

[24] Lindsey Power, Cédric Allain, Thomas Moreau, Alexandre Gramfort, and Timothy Bardouille. Using convolutional dictionary learning to detect task-related neuromagnetic transients and ageing trends in a large open-access dataset. NeuroImage, 267:119809, 2023.

[25] Carlos H Mendoza-Cardenas and Austin J Brockmeier. Shift-invariant waveform learning on epileptic ecog. In 2021 43rd Annual International Conference of the IEEE Engineering in Medicine & Biology Society (EMBC), pages 1136–1139. IEEE, 2021.

[26] Hinrich Schütze, Christopher D Manning, and Prabhakar Raghavan. Introduction to information retrieval, volume 39. Cambridge University Press Cambridge, 2008.

[27] Josef Sivic and Andrew Zisserman. Efficient visual search of videos cast as text retrieval. IEEE Transactions on Pattern Analysis and Machine Intelligence, 31(4): 591–606, 2008.

[28] Karen Sparck Jones. A statistical interpretation of term specificity and its application in retrieval. Journal of Documentation, 28(1): 11–21, 1972.

[29] L. S. Shapley. A value for n-person games. In Harold William Kuhn and Albert William Tucker, editors, Contributions to the Theory of Games, Volume II, chapter 17, pages 307–318. Princeton University Press, Princeton, 1953.

[30] Erik Štrumbelj and Igor Kononenko. Explaining prediction models and individual predictions with feature contributions. Knowledge and Information Systems, 41: 647–665, 2014.

[31] Scott M Lundberg and Su-In Lee. A unified approach to interpreting model predictions. Advances in Neural Information Processing Systems, 30, 2017.

[32] David J Kwiatkowski, Hongbing Zhang, Jennifer L Bandura, Kristina M Heiberger, Michael Glogauer, Nisreen el Hashemite, and Hiroaki Onda. A mouse model of TSC1 reveals sex-dependent lethality from liver hemangiomas, and up-regulation of p70s6 kinase activity in Tsc1 null cells. Human Molecular Genetics, 11(5): 525–534, 2002.

[33] Stephen Robertson. Understanding inverse document frequency: On theoretical arguments for IDF. J. Doc., 60(5): 503–520, 2004.

[34] E.P. Simoncelli, W.T. Freeman, E.H. Adelson, and D.J. Heeger. Shiftable multiscale transforms. IEEE Transactions on Information Theory, 38(2): 587–607, 1992.

[35] Ivana Tošić and Pascal Frossard. Dictionary learning. IEEE Signal Processing Magazine, 28(2): 27–38, 2011.

[36] Scott Shaobing Chen, David L Donoho, and Michael A Saunders. Atomic decomposition by basis pursuit. SIAM Review, 43(1): 129–159, 2001.

[37] S.G. Mallat and Zhifeng Zhang. Matching pursuits with time-frequency dictionaries. IEEE Transactions on Signal Processing, 41(12): 3397–3415, 1993.

[38] Yagyensh Chandra Pati, Ramin Rezaiifar, and Perinkulam Sambamurthy Krishnaprasad. Orthogonal matching pursuit: Recursive function approximation with applications to wavelet decomposition. In Proceedings of 27th Asilomar Conference on Signals, Systems and Computers, pages 40–44. IEEE, 1993.

[39] Joel A Tropp. Greed is good: Algorithmic results for sparse approximation. IEEE Transactions on Information theory, 50(10): 2231–2242, 2004.

[40] M. Aharon, M. Elad, and A. Bruckstein. K-svd: An algorithm for designing overcomplete dictionaries for sparse representation. IEEE Transactions on Signal Processing, 54(11): 4311–4322, 2006.

[41] Ron Rubinstein, Michael Zibulevsky, and Michael Elad. Efficient implementation of the k-svd algorithm using batch orthogonal matching pursuit. Technical Report CS-2008-08, Technion - Computer Science Department, 2008.

[42] Michal Aharon. Overcomplete dictionaries for sparse representation of signals. PhD thesis, Technion-Israel Institute of Technology, Faculty of Computer Science, 2006.

[43] J.J. Thiagarajan, K.N. Ramamurthy, and A. Spanias. Shift-invariant sparse representation of images using learned dictionaries. In IEEE Workshop on Machine Learning for Signal Processing, pages 145–150, Oct 2008.

[44] N Locantore, JS Marron, DG Simpson, N Tripoli, JT Zhang, KL Cohen, Graciela Boente, Ricardo Fraiman, Babette Brumback, Christophe Croux, et al. Robust principal component analysis for functional data. Test, 8: 1–73, 1999.

[45] Sangil Han, Sungkyu Jung, and Kyoowon Kim. Robust svd made easy: A fast and reliable algorithm for large-scale data analysis. In International Conference on Artificial Intelligence and Statistics, pages 1765–1773. PMLR, 2024.

[46] Wilson Truccolo, Kevin H Knuth, Ankoor Shah, Steven L Bressler, Charles E Schroeder, and Mingzhou Ding. Estimation of single-trial multicomponent erps: Differentially variable component analysis (dVCA). Biological cybernetics, 89(6): 426–438, 2003.

[47] Arthur E Hoerl and Robert W Kennard. Ridge regression: applications to nonorthogonal problems. Technometrics, 12(1): 69–82, 1970.

[48] Fabian Pedregosa, Gaël Varoquaux, Alexandre Gramfort, Vincent Michel, Bertrand Thirion, Olivier Grisel, Mathieu Blondel, Peter Prettenhofer, Ron Weiss, Vincent Dubourg, Jake Vanderplas, Alexandre Passos, David Cournapeau, Matthieu Brucher, Matthieu Perrot, Édouard Duchesnay, Fabian Pedregosa, Gaël Varoquaux, Alexandre Gramfort, Vincent Michel, Bertrand Thirion, Olivier Grisel, Mathieu Blondel, Peter Prettenhofer, Ron Weiss, Vincent Dubourg, Jake Vanderplas, Alexandre Passos, David Cournapeau, Matthieu Brucher, Matthieu Perrot, and Édouard Duchesnay. Scikit-learn: Machine Learning in Python. Journal of Machine Learning Research, 12: 2825–2830, 2011.

[49] Tom Fawcett. ROC graphs: Notes and practical considerations for researchers. Machine Learning, 31(1): 1–38, 2004.

[50] Jack Hogan and Niall M Adams. On averaging ROC curves. Transactions on Machine Learning Research, 2023.

[51] Mircea Steriade, Angel Nunez, and Florin Amzica. Intracellular analysis of relations between the slow (< 1 hz) neocortical oscillation and other sleep rhythms of the electroencephalogram. Journal of Neuroscience, 13(8): 3266–3283, 1993.

[52] P Achermann and AA Borbély. Low-frequency (< 1 hz) oscillations in the human sleep electroencephalogram. Neuroscience, 81(1): 213–222, 1997.

[53] Vincenzo Crunelli, Magor L Lőrincz, Adam C Errington, and Stuart W Hughes. Activity of cortical and thalamic neurons during the slow (< 1 Hz) rhythm in the mouse in vivo. Pflügers Archiv-European Journal of Physiology, 463(1): 73–88, 2012.

[54] L Veloso, J McHugh, Eva von Weltin, Sebas Lopez, I Obeid, and Joseph Picone. Big data resources for EEGs: Enabling deep learning research. In 2017 IEEE Signal Processing in Medicine and Biology Symposium (SPMB), pages 1–3. IEEE, 2017.

[55] Luca Pion-Tonachini, Ken Kreutz-Delgado, and Scott Makeig. ICLabel: An automated electroencephalographic independent component classifier, dataset, and website. Neuroimage, 198: 181–197, 2019.

[56] James Foulds and Eibe Frank. A review of multi-instance learning assumptions. The Knowledge Engineering Review, 25(1): 1–25, 2010.

[57] Thomas G Dietterich, Richard H Lathrop, and Tomás Lozano-Pérez. Solving the multiple instance problem with axis-parallel rectangles. Artificial Intelligence, 89(1–2): 31–71, 1997.

[58] Charles D Woody. Characterization of an adaptive filter for the analysis of variable latency neuroelectric signals. Medical and Biological Engineering, 5: 539–554, 1967.

